# The epigenetic factor *Zrf1* regulates intestinal stem cell proliferation during midgut regeneration

**DOI:** 10.1101/2025.01.16.633419

**Authors:** Joshua Shing Shun Li, Ying Liu, Ah-Ram Kim, Mujeeb Qadiri, Jun Xu, Baolong Xia, Richard Binari, John M. Asara, Yanhui Hu, Norbert Perrimon

**Affiliations:** Department of Genetics, Blavatnik Institute, Harvard Medical School, Boston, MA, USA; Department of Medicine, Harvard Medical School, Boston, MA, USA; Division of Signal Transduction, Beth Israel Deaconess Medical Center, Boston, MA, USA; Howard Hughes Medical Institute, Boston, MA, USA

## Abstract

Stem cells are essential for tissue maintenance and regeneration, balancing self-renewal and differentiation to support homeostasis and repair. Through an RNAi screen in the *Drosophila* midgut, we identified the epigenetic factor Zrf1 as a critical regulator of intestinal stem cell (ISC) proliferation. Functional analyses reveal that Zrf1 integrates inputs from multiple signaling pathways and interacts with components of the RNA-induced silencing complex (RISC). Zrf1 mutants exhibit elevated expression of transposable elements (TEs) and chromatin disruption, highlighting a broader role in genome stability. Single-nuclei RNA sequencing (snRNA-seq) further demonstrated the influence of Zrf1 on chromatin organization and TE repression, particularly within stem cell progenitors. Our findings suggest that Zrf1 is potentially a key chromatin regulator necessary for maintaining stem cell proliferation and genome integrity, enhancing our understanding of the molecular controls underlying stem cell function and chromatin dynamics.

## Introduction

Stem cells are fundamental to the maintenance and regeneration of tissues across multicellular organisms. These cells possess the remarkable ability to both self-renew and differentiate into various cell types, ensuring tissue homeostasis and repair. Understanding the molecular mechanisms that regulate stem cell behavior is crucial for advancing regenerative medicine and cancer therapy.

In recent years, the *Drosophila* midgut has emerged as a well-established system for studying mechanisms that maintain stemness and stem cell differentiation.^1, 2^ The fly midgut, a tube lined by an epithelial monolayer, is primarily composed of secretory enteroendocrine cells (EEs) and absorptive enterocytes (ECs). Resident intestinal stem cells (ISCs) continuously divide to replenish these cells as the midgut responds to damage. Many of the major pathways that orchestrate this process in *Drosophila* ISCs, including Notch, JAK/STAT, Hippo, Insulin, EGF and Wnt pathways, have also been shown to regulate mammalian epithelial stem cells. During periods of high proliferative pressure, ISCs are at risk of replication-induced DNA damage that can result in chromatin misregulation and uncontrolled retrotransposon expression.

Given the mutagenic potential and genomic instability caused by transposable elements (TE), stem cells have evolved mechanisms to regulate the expression of endogenous TEs. In particular, in germline stem cells, *Piwi* prevents TE expression.^3^ More recently, *Piwi* was also found to repress TE expression in the midgut progenitors.^4, 5^ While there is some disagreement in the phenotypic consequence of perturbing *Piwi*, both studies suggest the importance of maintaining proper heterochromatin organization in maintaining stem cell proliferation. Increasingly, other studies in *Drosophila* have also highlighted the role of chromatin regulators in ISC proliferation and differentiation.^6–8^

In a screen biased toward DNA binding proteins, we identified CG10565 as a regulator of ISC proliferation in the adult *Drosophila* midgut. *CG10565* (hereafter as *Zrf1*) encodes the mammalian ortholog, Zuotin-related factor 1 (ZRF1), which has been shown to displace polycomb-repressive complex 1 from chromatin to facilitate transcriptional activation.^2^ We found that Zrf1 is epistatic to multiple signaling pathways that regulate ISC proliferation and acts downstream of Myc, which has been shown to integrate proliferation cues from multiple signaling pathways.^9^ By leveraging genetic tools, immunoprecipitation-mass spectrometry (IP-MS), AlphaFold-Multimer (AFM) analyses, and single-nucleus RNA sequencing (snRNA-seq), we present a comprehensive understanding of the role of Zrf1 in ISC proliferation and chromatin regulation. Our findings contribute to the broader context of how chromatin regulators influence stem cell biology and tissue regeneration.

## Results

### *Zrf1* is necessary for ISC proliferation during homeostasis and regeneration

To identify intrinsic factors essential for ISC proliferation, we performed an RNAi screen of candidate genes biased towards DNA-binding proteins enriched in intestinal progenitors.^10–12^ Utilizing the *Escargot (esg)-Gal4* combined with the TARGET system (*tub-Gal80^ts^*),^13^ we selectively drove *UAS-RNAi* in adult ISC/EBs (*esg^ts^*). Progenitors were visualized using *UAS-GFP*, and anti-phospho-histone3 (pH3) staining served as a marker for ISC mitosis. Among ∼120 RNAi lines targeting ∼73 genes, three independent RNAi transgenes targeting *CG10565 (*hereafter referred to as *Zrf1*) significantly reduced ISC proliferation during homeostasis and in response to damage-inducing diets (Fig S1A, 1A-1C). *Zrf1* knockdown also resulted in fewer *Delta-LacZ*-positive and GFP-positive cells (Fig 1D, 1E, 1J and 1K). Lineage tracing using the *esg^ts^F/O* system^14^ revealed that progenitors failed to divide following *Zrf1 RNAi* knockdown (Fig 1F, 1G, 1J, S1B and S1C). CRISPR/Cas9-mediated knockout (*esg^ts^>Cas9.P2)* of *Zrf1* in ISC/EBs produced a similar phenotype (Fig 1H-1J). Importantly, the reduction in ISC proliferation was not a result of cell death since blocking apoptosis, by overexpressing either *DIAP* or *p35* in progenitors, could not rescue this defect (Fig 1C). The proliferation in response to damage in ISCs was unaffected when *Zrf1* was downregulated in adult enteroblasts (*Su(H)^ts^*) or EEs (*pros^ts^*) (Fig 1L). Regeneration of ISCs was only impaired when *Zrf1* was knocked down specifically in ISCs (*esg^ts^+Su(H)-Gal80* or *Dl^ts^*)(Fig 1L and S1E).

**Figure 1:**
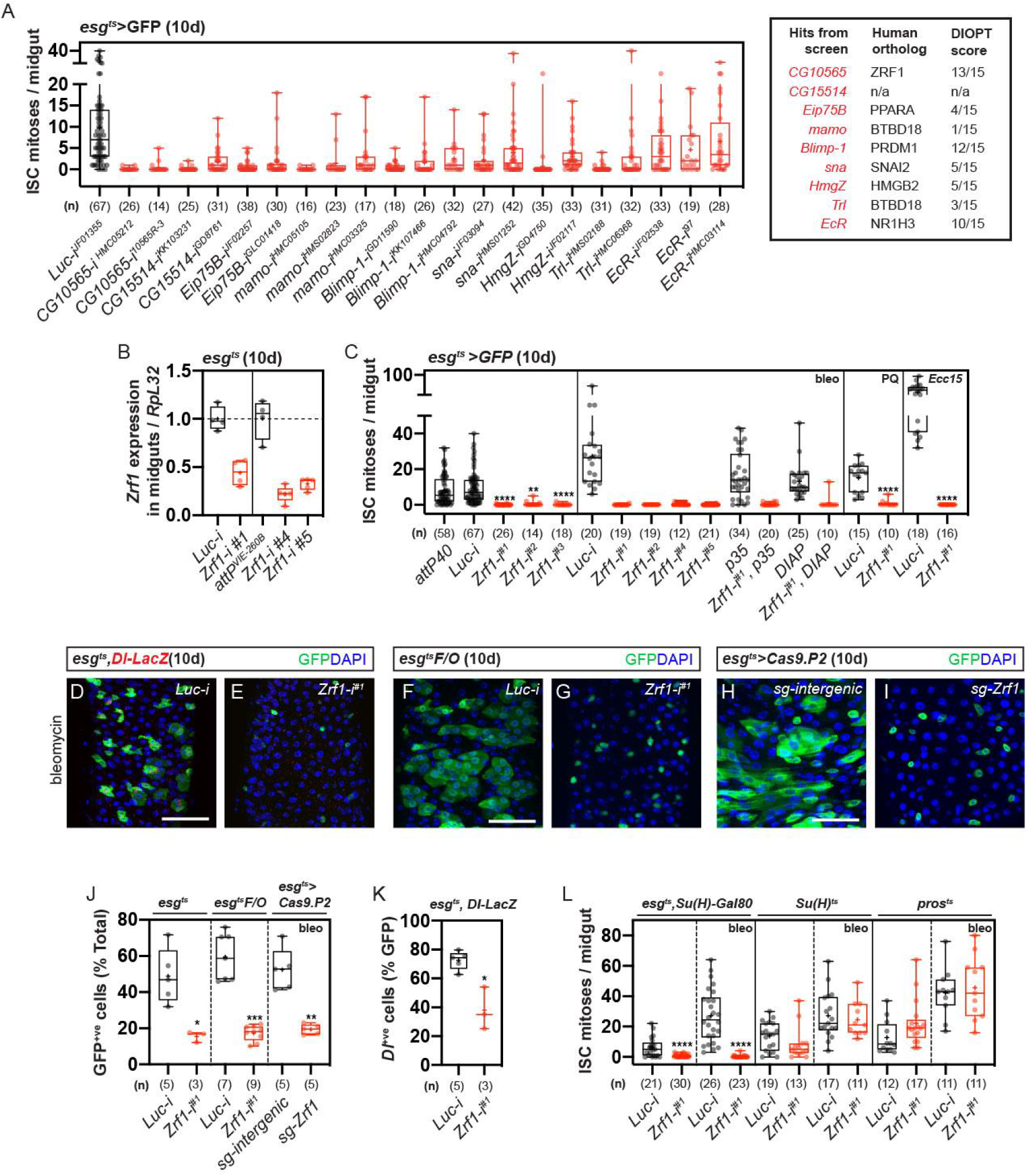
A screen identifies Zrf1 as a regulator of ISC proliferation during homeostasis and in response to damage. (A) Box plot showing ISC mitoses per midgut in *esg^ts^>GFP* flies after 10 days of RNAi induction targeting candidate genes enriched in intestinal progenitors. Hits from the screen are listed with their human orthologs and DIOPT scores. *Zrf1* RNAi lines (HMS00624, KK105682) led to reduced ISC mitosis compared to the control (*Luciferase, Luc*). Numbers below each box plot indicate the number of midguts analyzed. (B) Quantification of *Zrf1* transcript levels in midguts after 10 days of *Zrf1* knockdown in midgut progenitors, showing reduced expression relative to the housekeeping gene *RpL32* compared to controls. (C) Box plot showing ISC mitoses per midgut after *Zrf1* knockdown during homeostasis and following treatment with damage-inducing agents (bleomycin, paraquat, Ecc15). ISC mitoses are also shown following co-expression of anti-apoptotic factors (DIAP or p35). Numbers below each plot indicate the number of midguts analyzed. (D–H) Confocal images of midguts from *esg^ts^* flies treated with bleomycin. (D) Control (*Luciferase, Luc*). (E) *Zrf1* knockdown showing reduced ISC proliferation. (F– G) Lineage tracing using *esg^ts^F/O* system: (F) Control (*Luc*) and (G) *Zrf1* knockdown. (H) CRISPR/Cas9-mediated knockout of *Zrf1* (*esg^ts^>Cas9.P2*) showing similar reduction in ISC division. GFP marks progenitor cells; DAPI marks nuclei. Scale bar: 50 µm. (J) Box plot quantifying the percentage of GFP+ cells from experiments after *Zrf1* knockdown and control (Luc) in midgut progenitors. (K) Quantification of Dl-LacZ positive cells (ISC) after midgut-progenitor-specific *Zrf1* knockdown in comparison to control. (L) Box plots showing ISC mitoses per midgut after *Zrf1* downregulation in specific gut cell types: ISCs (*esg^ts^+Su(H)-Gal80*), enteroblasts (*Su(H)^ts^*) and enteroendocrine cells (*pros^ts^*). Midguts were treated with bleomycin. Numbers below each box plot indicate the number of midguts analyzed.

Altogether, these results indicate that *Zrf1* is cell-autonomously required for the proliferation of ISCs during homeostasis and regeneration.

### *Zrf1* promotes ISC proliferation in response to damage

To determine whether *Zrf1* is sufficient to promote ISC proliferation, we tested transgenes for effective *Zrf1* overexpression. We independently drove overexpression of four transgenes in ISC/EBs. These included two EP-like insertions, *P{EP}G4964* and *EP{EPgy2}Y14386*, which both consist of a *UAS* sequence inserted into the 5’ region upstream of the endogenous *Zrf1* gene as well as two *UAS-FlyORF* lines (*Zrf1^ORF.3xHA^* and *Zrf1^ORF.CC^*). Overexpression of all *Zrf1* constructs resulted in a 3-to-5-fold increase in *Zrf1* levels with the exception of *P{EP}G4964* (Fig 2A). Interestingly, upregulating *Zrf1* in progenitors only led to a moderate increase in ISC proliferation in response to damage but not during homeostasis (Fig 2B). Furthermore, CRISPR/Cas9-mediated transcriptional activation (*esg^ts^>Cas9.VPR)* of *Zrf1* also led to a damage-specific increase in ISC proliferation (Fig 2C and 2D). These results suggest that Zrf1 is sufficient to increase ISC proliferation only in response to damage.

**Figure 2:**
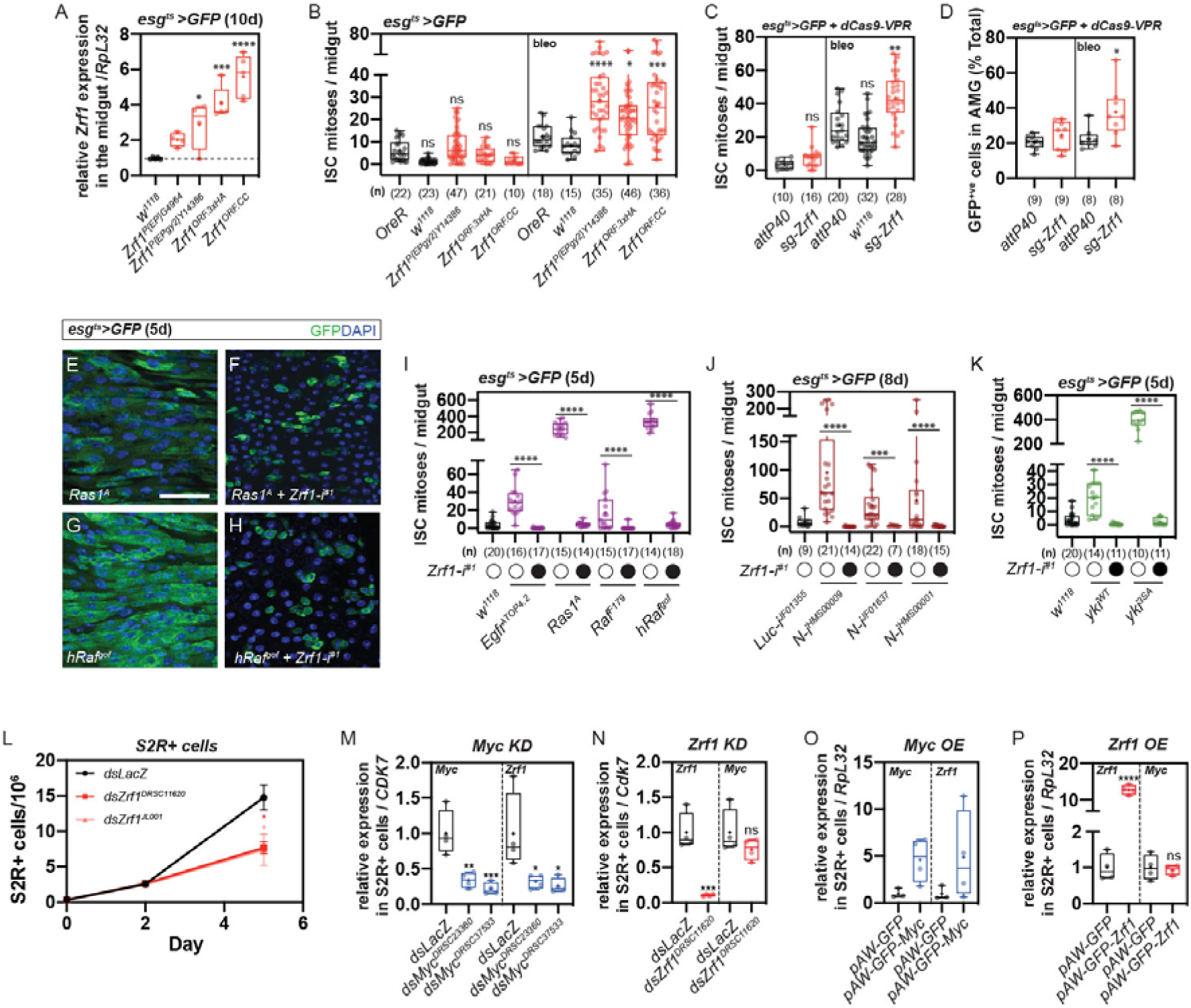
Zrf1 promotes ISC proliferation in response to damage. (A) Relative *Zrf1* expression levels in midguts from *esg^ts^>GFP* flies after 10 days of overexpression using different transgenes. Expression was measured relative to the housekeeping gene *RpL32*. Overexpression of *Zrf1* (EPgy2Y14386, ORF.3xHA, and ORF.CC) resulted in increased *Zrf1* transcript levels compared to the control (wild type, w^1118^). (B) ISC mitoses per midgut in *esg^ts^>GFP* flies overexpressing *Zrf1* transgenes (EPgy2Y14386, ORF.3xHA, ORF.CC) during homeostasis and after bleomycin treatment (bleo). Number of midguts analyzed is shown below each box plot. (C) Quantification of ISC mitoses in whole midguts of *esg^ts^>GFP* + *dCas9-VPR* flies, comparing *sg-Zrf1* to controls (*attP40, sg-intergenic*) under homeostasis and bleomycin treatment conditions. (D) Quantification of GFP+ progenitor cells in the anterior midgut (AMG) of *esg^ts^>GFP* + *dCas9-VPR* flies, showing the effect of sg-Zrf1 compared to sg-intergenic controls. (E–H) Confocal images of ISC/EB progenitors in *esg^ts^>GFP* flies overexpressing fly *Ras1A* or human gain-of-function Raf (*hRaf^gof^*) alone or in combination with *Zrf1* knockdown. Knockdown of *Zrf1* rescued ISC overproliferation induced by activated Ras1 and hRaf. GFP marks progenitor cells; DAPI marks nuclei. Scale bar: 50 µm. (I) ISC mitoses per midgut after 5 days of ectopic activation of the EGFR/Ras/Raf signaling components (Egfr^TOP4.2^, Ras1A, Raf^F149^, hRaf^gof^) in *esg^ts^>GFP* flies with and without *Zrf1* knockdown (Zrf1HMS00624). (J) ISC mitoses per midgut after 8 days of Notch knockdown in *esg^ts^>GFP* flies with and without *Zrf1* knockdown. (K) ISC mitoses per midgut after 5 days of overexpressing wildtype *Yorkie* (*Yki^WT^*) or constitutively active *Yorkie* (*Yki^3SA^*) in *esg^ts^>GFP* flies with and without *Zrf1* knockdown. (L) Growth curves of S2R+ cells over a 6-day period showing cell proliferation after dsRNA mediated knockdown of *Myc* (*dsMyc*), *Zrf1* (*dsZrf1*), and *LacZ*(*dsLacZ,* control). (M) Relative expression of *Myc* and *Zrf1* in S2R+ cells following knockdown of *Myc*, showing that *Myc* knockdown reduces *Zrf1* transcript levels but not vice versa. (N) Relative expression of *Myc* and *Zrf1* in S2R+ cells following knockdown of *Zrf1*, indicating that *Zrf1* knockdown does not affect Myc expression. (O) Relative expression of *Myc* and *Zrf1* following *Myc* overexpression (*pAWGFP-Myc*) in S2R+ cells, showing that *Myc* overexpression increases *Zrf1* transcript levels. (P) Relative expression of *Myc* and *Zrf1* in S2R+ cells following *Zrf1* overexpression (*pAWGFP-Zrf1*), showing that *Zrf1* overexpression does not affect *Myc* transcript levels.

### *Zrf1* is epistatic to multiple signaling pathways to regulate ISC proliferation

Bleomycin-induced damage in the gut is known to activate EGFR signaling.^15^ Since the loss-of-function and overexpression of *Zrf1* resulted in the most pronounced defects in response to damage, we sought to test if it is epistatic to the EGFR/Ras/Raf pathway. Ectopic expression of activated EGFR (*TOP4.2*), Ras (*Ras1^A^*), Raf (*Raf^F1^*^49^) or the human Raf (*hRaf^gof^*) in progenitors resulted in ISC overproliferation (Fig 2E, 2G and 2I), which could be rescued by *Zrf1* knockdown (Fig 2F, 2H and 2I), which places it downstream of the EGFR/Ras/Raf pathway.

However, *Zrf1* is not restricted to EGFR signaling, since epistatic experiments place it downstream of both Notch and Yorkie pathways as well (Fig 2J and 2K). These findings imply a broad function for *Zrf1*, reminiscent of the role of *Myc* in the midgut.^9^ A previous study showed that knockdown of *Myc* in ISCs caused stunted proliferation in response to damage, and like *Zrf1*, downregulating *Myc* rescued overproliferation defects caused by various signaling pathways.^9^ Thus, we suspected Zrf1 to be in the same pathway as Myc and found compelling evidence to support this when examining the literature. ChIP-seq of Myc in Kc cells showed that Myc binds to *Zrf1*.^16^ Microarrays of larval tissue indicate that *Zrf1* is upregulated following *Myc* overexpression.^17^ Finally, co-expression analysis of scRNA-seq data show *Zrf1* as a putative member of the *Myc* regulon.^18^

To test whether *Zrf1* is downstream of Myc, we knocked down *Myc* in S2R+ cells which resulted in a decrease in *Zrf1* transcript levels (Fig 2M). Knockdown of *Zrf1* stunted S2R+ proliferation but did not affect the levels of *Myc* (Fig 2L and 2N).

Accordingly, *Myc* overexpression increased the levels of *Zrf1*, whereas upregulating *Zrf1* did not affect *Myc* levels (Fig 2O and 2P). In conclusion, our results suggest that *Zrf1* could function downstream of *Myc*, integrating multiple signaling pathways to regulate ISC proliferation.

### Zrf1 Protein-Protein interaction network identifies association with RISC

To gain insight into the function of Zrf1, we performed a search for interacting proteins, which we reasoned would identify regulators and localization partners. The Zrf1 protein complex was immunoprecipitated under mild detergent conditions (NP40) using GFP-trap beads from S2R+ cells expressing N-terminally GFP-tagged Zrf1 and subjected to mass spectrometry (IP-MS) (Fig 3A, 2B and S2A-S2C). Since the IP-MS dataset contains a mixture of direct and indirect protein-protein complexes, we employed sequential AlphaFold-Multimer (AFM) screening to predict direct protein-protein interactions, applying Local Interaction Score (LIS) and Local Interaction Area (LIA).^19^ (Fig 3C) First, we predicted the direct interactors with Zrf1, which also interact with each other within this group. To further expand the PPI network, we conducted second and third rounds of AFM screening, resulting in over 500 predictions. Ultimately, the Zrf1 PPI network comprising approximately 40 direct PPIs was built. The details for whole prediction results and predicted aligned error maps for all positive PPI predictions are shown in Figure S3. Network analysis revealed associations with ribosomal proteins and components of the RNA-induced silencing complex (RISC) among other proteins (Fig 3C). Consistent with this, Zrf1 IP-MS showed >2-fold spectral counts for Rm62, Fmr1, vig, vig2, AGO2 and Tdrd3 in comparison to controls. To test whether these interacting proteins associated with Zrf1, we conducted co-immunoprecipitation (Co-IP) experiments with S2R+ cells transfected with C-terminally 3xHA-tagged Zrf1 and N-terminal-GFP-tagged binding partners (Fig 3D). In total, Zrf1-3xHA immunoprecipitated with 17 out of 19 GFP-tagged binding partners. This included CG7182, CG8635, Hsc70-4, mod, penguin, rin, Rm62, vig, two isoforms of vig2, FDY (a Y-chromosome duplication of *vig2*), CG8545, 128up and Tdrd3. Reciprocal Co-IP found that C-terminal-3xHA-tagged Rm62, Fmr1, vig, vig2a, vig2b, AGO2, FDY and Tdrd3 precipitated with GFP-Zrf1 (Fig 3E and S4A-S4D). Antibodies against these proteins showed that endogenous levels also immunoprecipitate with Zrf1, indicating a robust physical association between Zrf1 and RISC components (Fig S4E and 3F).

**Figure 3:**
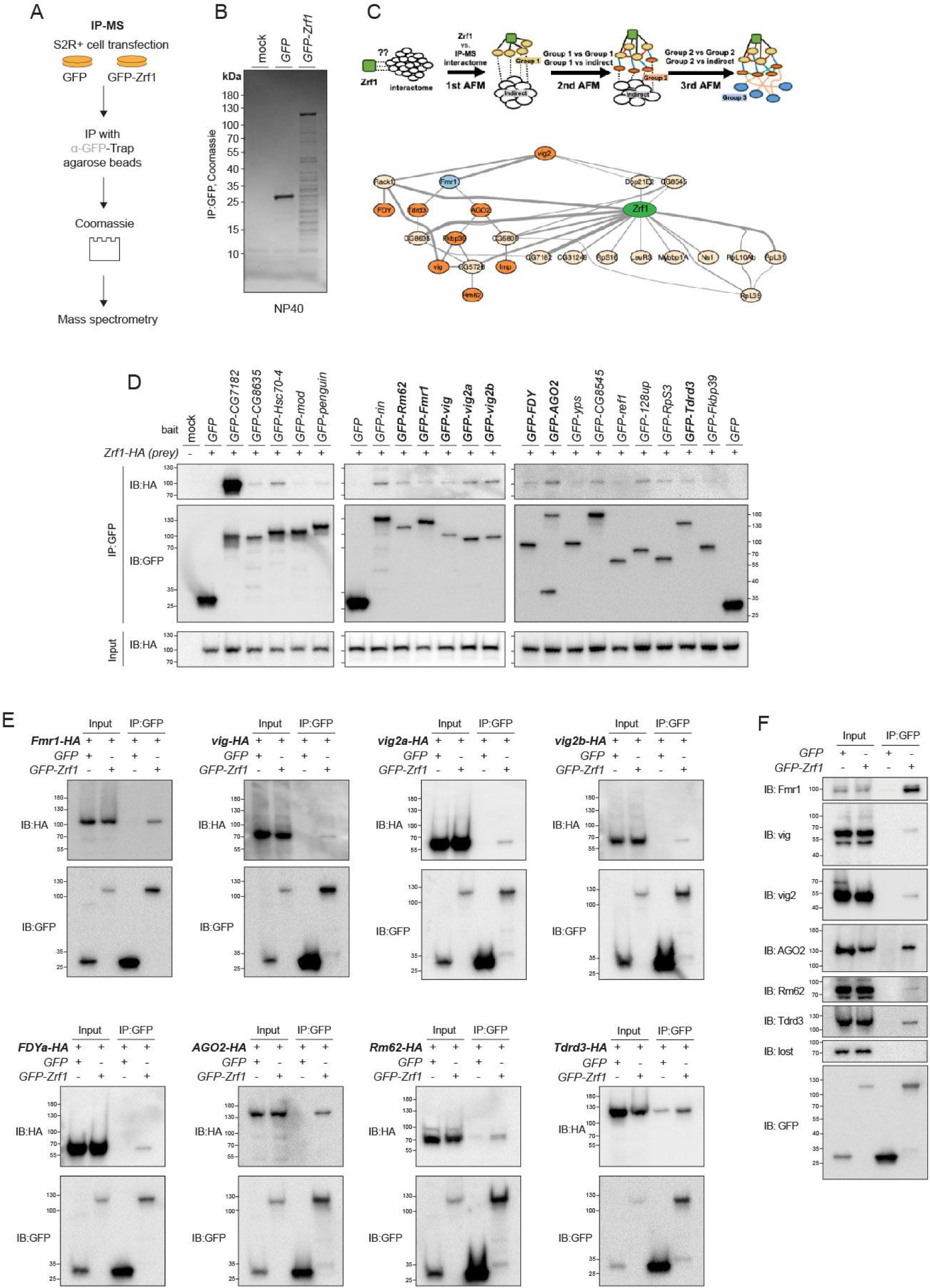
IP-MS-AFM reveals Zrf1 Protein-Protein Interactions (PPI) with Components of RISC in *Drosophila*. (A) Schematic of the experimental workflow for immunoprecipitation-mass spectrometry (IP-MS). S2R+ cells were transfected with constructs expressing either GFP or GFP-Zrf1. Following cell lysis, GFP-trap agarose beads were used to immunoprecipitate complexes, which were then analyzed via mass spectrometry after Coomassie staining. (B) Coomassie-stained gel showing proteins immunoprecipitated with GFP-Zrf1 compared to GFP alone. (C) Sequential AlphaFold-Multimer (AFM) screen scheme and protein interaction network generated from the Zrf1 IP-MS interactome analysis. The screening begins with the 1st AFM, predicting interactions between Zrf1 (central node, green) and the entire interactome to identify direct interactors with Zrf1 (Group 1, beige nodes). The 2nd AFM further explores interactions within Group 1 and between Group 1 and the indirect interactome, identifying new interactors (Group 2, orange nodes) and connections within Group 1. Finally, the 3rd AFM involves predictions between Group 2 proteins and additional indirect interactors, forming Group 3 (light blue nodes). Edge thickness represents the Local Interaction Score (LIS), where thicker lines indicate stronger predicted interactions. (D) Validation of Zrf1 interactions with candidate proteins identified in IP-MS through co-immunoprecipitation (Co-IP). S2R+ cells were transfected with GFP-tagged candidate proteins and HA-tagged Zrf1. Immunoprecipitation with anti-GFP was followed by immunoblotting with anti-HA to detect Zrf1. Representative interactions with various proteins, including Rm62, vig, and others, are shown. (E) Reciprocal Co-IP with GFP-tagged Zrf1 and HA-tagged components of RISC, such as Fmr1, vig, vig2a, and others. (F) Immunoblots validating the association of Zrf1 with endogenous RISC components. Proteins including Fmr1, vig, vig2, AGO2, Rm62, and Tdrd3 were detected, confirming the interactions of Zrf1 with the RISC complex.

To determine if the physical interactions between zrf1 and RISC was conserved in humans, we conducted IP-MS on N- and C-terminally epitope-tagged ZRF1 in HEK293T cells (under mild detergent conditions) (Fig S5A-S5C). The MS spectrum counts of both tagged versions of ZRF1 enriched for SERBP1 (ortholog of vig, vig2 & FDY), human orthologs of Rm62 (DDX17, DDX21 and DDX5) or Fmr1 (FXR1, FMRP and FXR2) but not for TDRD3, HABP4 or members of the AGO-family (Fig S5D).

Nonetheless, we conducted Co-IP experiments to determine if the PPIs of Zrf1 is conserved. C-terminal-epitope-tagged SERBP1, HABP4, DDX17, DDX21, or DDX5 all precipitated with Flag-ZRF1 (Fig S5F top). ZRF1-GFP precipitated with N-terminal-Flag-tagged AGO1 (Fig S5F bottom). Reciprocal Co-IP confirmed the ZRF1-SERBP1 and ZRF1-HABP4 interactions. Additionally, IP with a ZRF1 antibody successfully detected endogenous FXR1, FMRP and FXR2 (Fig S5E).

### Components of RISC genetically interact with *Zrf1* to regulate ISC proliferation

We hypothesized that perturbing components of RISC would mimic the effects of *Zrf1* knockdown. Utilizing CRISPR, we generated mutants for *vig* and *vig2*, and assessed ISC proliferation during gut damage (Fig 4A). Homozygous and transheterozygous mutants of *vig*, *vig2*, and *AGO2* exhibited reduced ISC proliferation in response to bleomycin, while FDY mutants showed no defects (Fig 4B). Additionally, double heterozygous animals — *Zrf1^2^*^42^/+ with either fmr1(*fmr1^d113M^/Zrf1^2^*^42^), vig2 (vig2*^Gd^*^59–75^/*Zrf1^2^*^42^), or *AGO2* (*AGO2/+;Zrf1^2^*^42^*/+*) — displayed a synergistic decrease in ISC proliferation compared to individual heterozygotes (Fig 4C). These results collectively suggest a genetic interaction between *Zrf1* and the RISC components (*Fmr1*, *vig2*, and *AGO2*) pivotal for regulating ISC proliferation during damage recovery.

**Figure 4:**
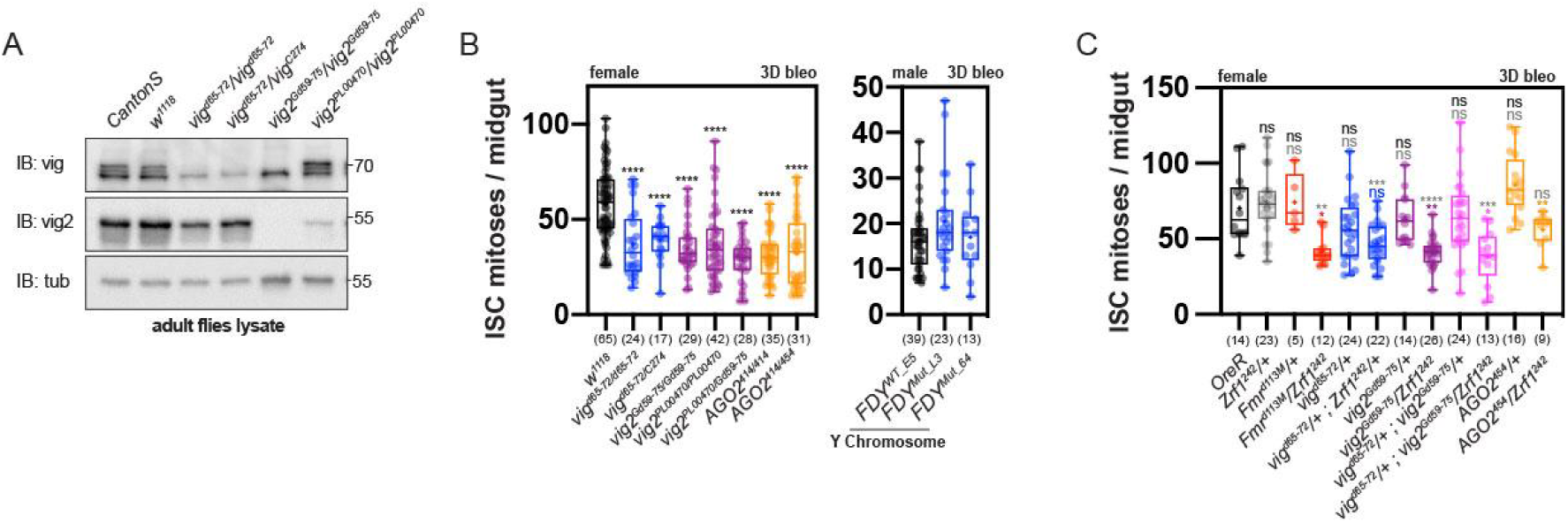
Genetic Interaction Between Zrf1 and RISC Components Regulates ISC Proliferation during midgut damage. (A) Western blot analysis of adult fly lysates showing protein levels of Vig and Vig2 in various genetic backgrounds. Samples include wild-type controls (*Canton-S, w^1118^*), and mutant alleles for *vig* (*vig^d^*^65–72^*, vig^d^*^63–73^*, vig^C274^*), *vig2* (*vig2^Gd^*^59–75^*, vig2^PL00470^*), and a double mutant (*vig2^PL00470^/vig2^Gd^*^59–75^). Tubulin (tub) was used as a loading control. The blots confirm the effective reduction of Vig and Vig2 in the mutant backgrounds. (B) ISC mitoses per midgut after 3 days of bleomycin (bleo) treatment in various female and male genotypes. Mutants for *vig* and *vig2* exhibit a reduction in ISC proliferation compared to controls (*w^1118^*). Mutants for *AGO2* (*AGO2^414/414^* and *AGO2^414/454^*) also show decreased proliferation. (C) ISC mitoses per midgut after 3 days of bleomycin treatment in double heterozygous animals. Females carrying *Zrf1^2^*^42^*/+* combined with heterozygous mutant alleles for RISC components (*Fmr1^d113M^, vig^d^*^65–72^*, vig^d^*^63–73^*, vig2^Gd^*^59–75^, and *AGO2^454^*) are compared to individual heterozygotes. Significant reductions in ISC proliferation are observed in certain double heterozygotes, suggesting a synergistic interaction between Zrf1 and these RISC components. Error bars represent the interquartile range; n = number of midguts analyzed for each genotype. Box plots show median, quartiles, and range. Statistical significance determined by Student’s t-test; ns = not significant, **p < 0.01, ****p < 0.0001.

### *Zrf1* is not involved in RNAi-mediated post-transcriptional silencing

In light of the established roles of *AGO2* and *Rm62* in RNAi-mediated post-transcriptional silencing^20, 21^ and the physical association between Zrf1 and RISC, we posited that Zrf1 might play a role in the RNAi pathway. To investigate, we assessed the knockdown efficiency of *dsGFP* in S2R+ cells expressing GFP following pre-treatment with various dsRNA. Initially, cells were exposed to either *dsLacZ* (control), *dsAGO2*, or *dsZrf1*. After 3-5 days, the same cells were subjected to GFP dsRNA treatment or left untreated. The effectiveness of RNAi-mediated silencing was assayed by measuring GFP levels normalized to tubulin. As anticipated, RNAi-mediated GFP knockdown was compromised in cells pre-treated with *dsAGO2* compared to the control. Intriguingly, knockdown of *Zrf1* using two independent dsRNAs did not impact GFP knockdown (Fig 5A and 5B). Our findings indicate that *Zrf1* does not participate in RNAi-mediated post-transcriptional silencing.

**Figure 5:**
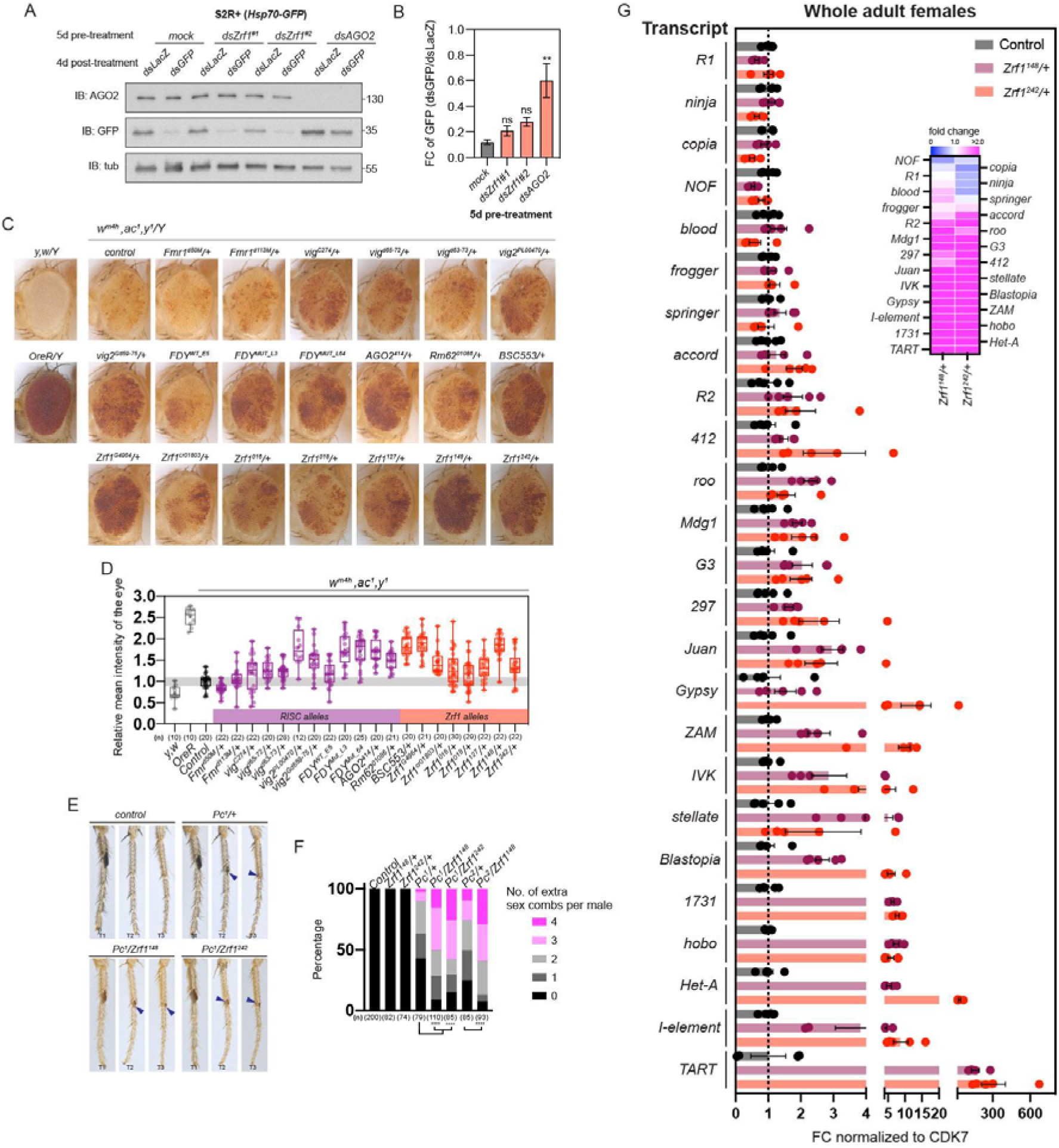
Zrf1 Mutants Exhibit Deregulation of Transposable Elements and Chromatin-Associated Defects. (A) Western blot analysis of S2R+ cells pre-treated for 5 days with various dsRNAs (*dsLacZ, dsZrf1, dsAGO2*). Cells were subsequently treated with *dsGFP* to assess RNAi-mediated silencing. Immunoblots show levels of AGO2, GFP, and tubulin (loading control). (B) Quantification of GFP levels from (A), normalized to tubulin. Knockdown of *Zrf1* did not significantly change GFP silencing efficiency, supporting that Zrf1 is not involved in RNAi-mediated post-transcriptional silencing. (C) Representative images of fly eyes of *w^m4h^, ac^1^/ y* males crossed with various RISC and *Zrf1* mutants. (D) Quantitative analysis of relative mean pigmentation intensity from (C). Box plots compare pigmentation across RISC and *Zrf1* mutant backgrounds, normalized to control (*w^m4h^, ac^1^/y*). Reduced pigmentation corresponds to enhanced suppression of PEV, while increased pigmentation indicates loss of PEV suppression. (E) Extra sex comb phenotypes in *Pc*^1^ and double heterozygous *Pc^1^; Zrf1* mutant males. Images show T2 and T3 legs of control (*Pc^1^/+*) and double heterozygous (*Pc^1^/Zrf1^148^, Pc^1^/Zrf1^127^*) flies. Blue arrows point to the presence of ectopic sex combs, indicating synergistic enhancement of *Pc*^1^ phenotypes by *Zrf1* mutants. (F) Percentage distribution of extra sex combs in *Pc*^1^ and double heterozygous *Pc^1^; Zrf1* mutant males. Bar graph displays the proportion of males with 0, 1, 2, 3, or 4 extra sex combs per male across genotypes, with *Zrf1* mutants showing an increased frequency of ectopic sex combs. (G) Fold change (FC) in transposable element (TE) expression levels in whole adult females from *Zrf1* mutant backgrounds (*Zrf1^148^/+* and *Zrf1^127^/+*) compared to control. FC values are normalized to *CDK7*. Dot plot and heatmap display upregulated TEs, with specific TEs showing significant deregulation. Heatmap inset indicates fold change values for a subset of upregulated TEs, demonstrating consistent patterns of TE misregulation in *Zrf1* mutants. Box plots display median, quartiles, and range; statistical significance determined by Student’s t-test; ns = not significant, **p < 0.01, ***p < 0.001, ****p < 0.0001.

**Figure 6.**
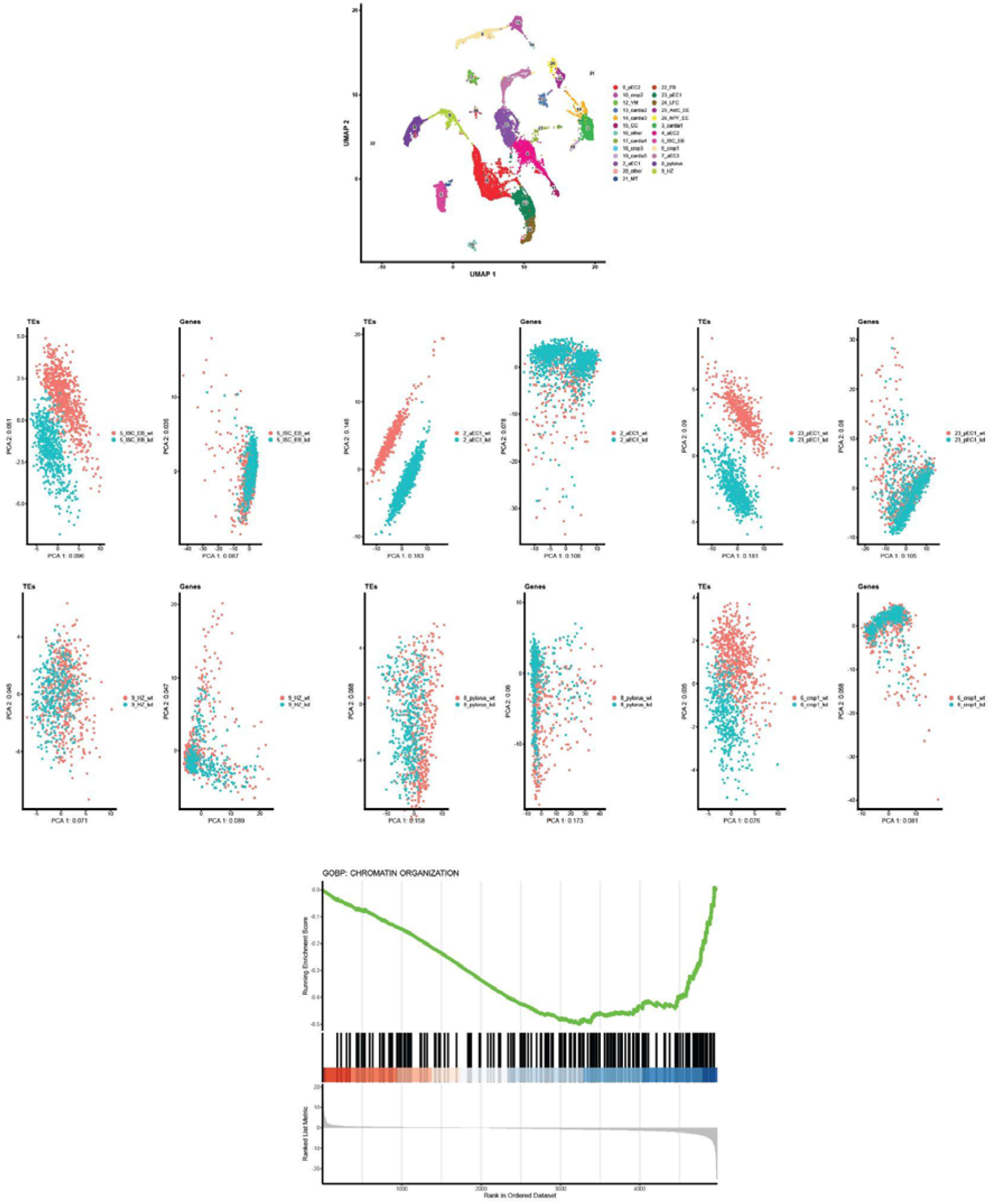
Single-nuclei and principal component analyses of gene and transposable element expression in *Zrf1* knockdown and control guts. (A) UMAP of single-nuclei RNA sequencing data from midgut cell types, showing clustering of different cell populations labeled by their cell type. (B-D) PCA plots showing separation of *Zrf1* knockdown *(Zrf1-i)* and control *(Luc-i*) samples within specific clusters based on either transposable elements (TEs) or gene expression. Each plot represents a specific cell type: ISC, aEC, pEC, and other midgut clusters. (E) GSEA plot for chromatin organization gene set enrichment in *Zrf1* knockdown.

### *Zrf1* mutants dominantly suppress position-effect variegation and enhance polycomb defects

Previous studies have revealed that RISC components have non canonical roles in modifying heterochromatin gene silencing.^22, 23^ To investigate whether Zrf1 shares similar functions, we used the positive-effect variegation (PEV) reporter *In(1)w^m4h^*, where the *white* (*w*) gene is transposed close to pericentromeric heterochromatin on the X-chromosome by a large chromosome inversion.^24^ PEV results in suppressed *w^+^* expression in *w^m4h^/Y* males, leading to mottled eye pigmentation (Fig 5C). Conventionally, enhancers and suppressors of PEV have been associated with genes that regulate chromatin condensing and de-condensing, respectively. Confirming earlier studies,^25, 26^ *AGO2* (*AGO2^414^/+*) and Rm62 (*Rm62^01086^/+*) mutants suppressed *w^m4h^* PEV, while *Fmr1* (*Fmr1^d50M^/+* & *Fmr1^d113M^/+*) mutations did not. In addition, *vig* alleles *(vig^d^*^65–72^*/+* & *vig^d^*^63–73^*/+* but not *vig^C274^/+*) moderately suppressed PEV, whereas *vig2* alleles (*vig2^PL00470^* & *vig2^Gd^*^59–75^) exhibited strong suppression. Finally, *FDY* mutations (*FDY^MUT_L3^* & *FDY^MUT_L64^*) also dominantly suppressed PEV when compared to the control (*FDY^WT_E5^*) (Fig 5C and 5D).

Similar to the components of RISC, heterozygous alleles of *Zrf1* dominantly suppressed *w^m4h^* PEV. This encompassed a deficiency that covers *Zrf1* (*BSC553*), a P-element insertion with a weak mini-white (*Zrf1^G4964^*) and five indel mutants (*Zrf1^016^, Zrf1^018^, Zrf1^127^, Zrf1^148^* & *Zrf1^2^*^42^) generated using CRISPR/Cas9 (Fig 5C, 5D and S6A). Collectively, our findings unveil Zrf1 and RISC components (except Fmr1) as dominant suppressors of PEV, suggesting a broad role in chromatin condensation.

Notably, Polycomb group proteins are also known to regulate transcriptional repression of target genes through chromatin modifications. A previous study highlighted the ability of ZRF1to b ind to H2Aub1 and displace PRC1, activating previously repressed genes.^27^ Another study also showed that plant ZRF1 possesses both PRC1-related and PRC1-unrelated functions.^28^ Given this context, we hypothesized that Zrf1 might interact with Polycomb (Pc), influencing chromatin structure and gene expression. Using *Pc*^1^ animals, a weak amorphic allele of *Pc*, we tested for genetic interactions between *Zrf1* and *Pc.* In wildtype flies, sex combs are only found at the distal end of the male foreleg (T1) (Fig 5E and S6B). However, heterozygous *Pc*^1^ males have varying degrees of extra sex combs in the second (T2) and third (T3) legs. *Zrf1* heterozygous mutant males (*Zrf1^148^/+, Zrf1^2^*^42^*/+*) are indistinguishable from control animals. Interestingly, double heterozygous animals of *Zrf1* and *Pc* alleles (*Zrf1^148^/Pc^1^, Zrf1^2^*^42^*/Pc*^1^) synergistically enhanced the sex comb defect observed in *Pc*^1^ heterozygous animals alone (Fig 5E, 5F and S6B), suggesting that Zrf1 enhances the de-repressed genes regulated by Pc.

### Retrotransposable element expression are deregulated in Zrf1 mutants

Given that chromatin-associated regulators often influence retrotransposable element (TE) expression,^29–31^ we explored this possibility in the context of Zrf1. Initial qPCR analysis of whole animals with *Zrf1* heterozygous mutants revealed increased expression of several TEs, suggesting that Zrf1 normally represses these elements (Fig 5G). This observation was consistent when qPCR was performed on midguts, confirming that Zrf1 plays a role in TE repression in this tissue.

To understand the combined effects of Zrf1 and Pc on TE expression, we performed qPCR to assess TE expression in *Zrf1* and *Pc* double mutants, and found that some TEs also exhibited synergistic and additive effects of derepression in the *Zrf1* and *Pc* double mutants compared to the single heterozygous mutants. This suggests that Zrf1 and Pc cooperate to repress TE expression, and that their interaction is crucial for maintaining chromatin state and genomic stability (Fig S7).

To further validate the role of Zrf1 in transposable element (TE) regulation and chromatin organization in intestinal stem cell (ISC)-derived lineages, we performed single nuclei RNA-seq (snRNA-seq) on adult midgut samples, comparing control (*egt>luc-i*) and *Zrf1* knockdown (*egt>zrf1-i*) conditions. Following unsupervised clustering we annotated a total of 21 clusters which overtly mapped to cell types/sub-types similar to previous single-cell sequencing studies conducted in the fly midgut.^8, 11, 18, 32, 33^ Next, we focused our analysis on three clusters affected by *esg^ts^* manipulation representing ISC/EB, anterior enterocytes (aEC), and posterior enterocytes (pEC). Three non-ISC-derived lineages were also used as a negative control (pyloris, crop and hybrid zone). Interestingly, principal component analysis (PCA) based solely on TE expression revealed distinct separation between *Zrf1* knockdown *(Zrf1-i)* and control *(Luc-i*) in ISC, aEC, and pEC clusters. This separation was not observed when PCA was conducted using gene expression alone, suggesting that Zrf1 specifically affects TE expression in ISC-derived lineages. In contrast, clusters from non-ISC-derived lineages displayed no separation between knockdown and control in both TE and gene expression analyses, further underscoring the ISC-specific impact of Zrf1 on TE regulation.

Additionally, Gene Set Enrichment Analysis (GSEA) for chromatin organization genes indicated a significant enrichment in Zrf1 knockdown conditions, supporting a crucial role for Zrf1 in chromatin dynamics within ISC progenitor cells.

## Discussion

Our study has uncovered a critical role for Zrf1 in regulating intestinal stem cell (ISC) proliferation and chromatin dynamics in the *Drosophila* midgut. By leveraging a combination of genetic tools, IP-MS, AlphaFold-Multimer predictions, and single-nucleus RNA sequencing (snRNA-seq), we provide comprehensive insights into the multifaceted functions of Zrf1. Here, we discuss the implications of our findings and propose potential avenues for future research and therapeutic applications.

### Zrf1 as a key regulator of ISC proliferation

Although *Zrf1* orthologs share some conserved roles across species, their functions have diverged considerably. In mammals, ZRF1 displaces polycomb complexes to activate developmental genes critical for cell fate transition.^27^ Similarly, in plant, *Zrf1* orthologs promote the activation of genes associated with flowering and growth.^28, 34^ By contrast, the yeast ortholog Zuotin operates as a molecular chaperone within the ribosome-associated complex, assisting nascent polypeptide folding and maintain proteostasis.^35^ The functional differences highlight the unique value of *Drosophila* as a model to uncover aspects of Zrf1 biology that may be obscured in other organisms.

Our results establish that Zrf1 is essential for ISC proliferation during both homeostasis and regeneration. Our RNAi screen identified Zrf1 as a critical factor, with its knockdown leading to a significant reduction in ISC proliferation. This phenotype was not rescued by blocking apoptosis, indicating a specific requirement for Zrf1 in ISC proliferation rather than cell survival. Interestingly, Zrf1 overexpression promoted ISC proliferation only in response to damage, suggesting that its role is tightly regulated by cellular context and environmental cues.

The integration of Zrf1 into multiple signaling pathways, including the EGFR/Ras/Raf, Notch, and Yorkie pathways, underscores its broad regulatory capacity. Our epistasis experiments position Zrf1 downstream of these pathways, akin to the role of Myc in the midgut. This finding aligns with previous studies showing that Myc and Zrf1 share common regulatory networks,^9^ further supported by ChIP-seq and microarray data.^16, 17^ Altogether, our results suggest that Myc likely regulates the direct transcription of Zrf1.

### Zrf1 protein-protein interaction network and association with RISC

The construction of a Zrf1 protein-protein interaction network using AlphaFold-Multimer and Local Interaction Score (LIS) analysis revealed significant associations with components of the RNA-induced silencing complex (RISC).^19^ Co-IP experiments confirmed these interactions, highlighting the potential involvement of Zrf1 in RNA processing and silencing mechanisms. However, our functional assays demonstrated that Zrf1 does not participate in RNAi-mediated post-transcriptional silencing, redirecting our focus toward its chromatin-related functions.

### Zrf1 as an epigenetic factor

Our findings that *Zrf1* mutants dominantly suppress position-effect variegation (PEV) and enhance Polycomb (Pc) defects provide compelling evidence for a role of Zrf1 in epigenetic regulation. The synergistic interaction between *Zrf1* and *Pc*^1^ alleles, resulting in enhanced sex comb phenotypes, suggests that Zrf1 modulates chromatin structure and gene expression in concert with Pc group proteins. This is consistent with previous reports indicating the ability of Zrf1 to displace Pc complexes from chromatin to activate gene expression.^27^

### Zrf1 and retrotransposable elements

Zrf1 affects the regulation of retrotransposable elements (TEs). qPCR analysis revealed increased TE expression in *Zrf1* heterozygous mutants, indicating a repressive role for Zrf1 on these elements. The derepression of TEs was further exacerbated in *Zrf1* and *Pc* double mutants, demonstrating synergistic and additive effects. This reinforces the idea that Zrf1 and Pc collaborate to maintain genomic stability by repressing TEs.

### Implications for cancer and therapeutic potential

The role of Zrf1 in regulating ISC proliferation and chromatin dynamics suggests potential implications for cancer biology. Given that Zrf1 can function as both a tumor suppressor and an oncogene depending on the context, understanding its regulatory networks could provide new targets for cancer therapy.^36, 37^ For example, in the context of gastric carcinoma, where Zrf1 is overexpressed,^38^ targeting its interaction with Pc and RISC components could offer novel therapeutic strategies.

Furthermore, the specificity of the effects of Zrf1on ISC and progenitor cells, as revealed by our snRNA-seq data, highlights the importance of cell-type-specific approaches in therapeutic interventions. The distinct separation of clusters in PCA plots based on TE expression suggests that Zrf1 could be targeted to modulate TE activity and chromatin states in specific cell populations, potentially reducing off-target effects and enhancing treatment efficacy.

### Future directions

Future research should aim to elucidate the precise molecular mechanisms underlying Zrf1’s interactions with Pc and RISC components. Additionally, investigating Zrf1’s role in other tissues and cancer types will be crucial for translating these findings into clinical applications. Understanding how Zrf1 integrates multiple signaling pathways to regulate stem cell behavior and chromatin dynamics will provide valuable insights into stem cell biology and open new avenues for therapeutic development.

In conclusion, our study highlights the critical role of Zrf1 in ISC proliferation and chromatin regulation, with significant implications for stem cell biology and cancer therapy. By unraveling the complex networks involving Zrf1, this study paves the way for future research and potential clinical applications targeting Zrf1 and its associated pathways.

## Methods

### RNAi screening

We used *esg-Gal4, UAS-GFP, tub-Gal80^ts^* (*esg^ts^*) to knockdown genes in adult ISCs/EBs and quantified ISC proliferation by counting the number of mitotic (pH3-positive) cells per midgut. *UAS-RNAi* lines were obtained from Bloomington (BDSC), Vienna (VDRC) and NIG/Kyoto *Drosophila* stock centers.

*UAS-RNAi* stocks that are homozygous lethal on second chromosomes were placed over *CyO, twi-Gal4, UAS-2EGFP* balancers. Virgin *esg^ts^* females were crossed to *UAS-RNAi* males and reared at 18°C (restrictive temperature). After eclosion, *esg^ts^* + *UAS-RNAi* females were kept at 18°C for 3 days and shifted to 29°C (permissive temperature) for 10 days before their whole midguts were dissected. Flies were transferred to fresh vials every 2-3 days. Mated females were used in all experiments and reared with male flies at the ratio of 2:1. Whole midguts were analyzed for pH3 positive counts.

### Infection and compound feeding

The OD_600_ of an overnight suspension of *Ecc15* was determined by measuring two diluted (1:10 & 1:100) aliquots of the culture. After centrifuging the culture at 3000g for 20min, the pellet was resuspended in 5% sucrose with a volume to a final OD_600_ of 100. Paper towels cut out to the shape of the vial base were soaked with 500uL of the concentrated *Ecc15* solution. For oral infection, flies were starved for 2h at 29°C in an empty vial before transferring to vials containing *Ecc15*.

### Immunohistochemistry and imaging

Whole midguts were dissected in PBS and fixed in 4% PFA in PBS at room temperature for 30 minutes. Fixed guts were washed in 0.1% Triton X-100 in PBS (PBST) once then blocked with a blocking buffer (0.1% Triton, 5%NDS in PBS) for 30 minutes at RT. Primary antibodies were incubated overnight at 4°C in the blocking buffer. Guts were washed 3x in a blocking buffer and incubated with secondary antibodies overnight at 4°C along with DAPI (1:1000 of 1mg/ml stock). After antibody staining, guts were washed 3x in PBST and mounted in Vectashield antifade mounting medium (Vector Laboratories H-1200).

Tape was used as a spacer to prevent coverslips from crushing the guts. Antibody dilutions used were as follows: rabbit anti-pH3 (1:1000, Millipore 06-570), mouse anti-Dachshund (1:20, DSHB AB_579773), chicken anti-GFP (1:1000, Abcam ab13970), donkey anti-rabbit 565 (1:2000, Molecular Probes A31572), goat anti-mouse 633 (1:2000, Thermo Scientific A-21240) and goat anti-chicken 488 (1:2000, Thermo Fisher Scientific A-11039). Guts were imaged on a spinning-disk confocal system, consisting of a Nikon Ti2 inverted microscope equipped with a W1 spinning disk head (50um pinhole, Yokogawa Corporation of America) and a Zyla 4.2 Plus sCMOS monochrome camera (Andor).

### RNA extraction and quantitative real-time PCR

10-15 midguts were dissected in 1xPBS and homogenized in 300uL of TRIzol (ThermoFischer, 15596-026) using RNase-free pestles. RNA was extracted using Zymo Direct-zol RNA MicroPrep kit (R2050) and subsequently DNase-treated using Turbo DNA free (AM1907). 400-450ng of the resulting RNA was reverse transcribed using Bio-Rad iScript Select cDNA synthesis kit (708896) and Sybr green (1708880) based qRT-PCR was performed to determine the levels of gene expression. qRT-PCR primers were designed using FlyPrimerBank.^39^ Primers that were not from the literature were tested for their efficiency by running qRT-PCR on serial dilutions of pooled cDNA. Only primers with efficiencies of greater than 85% were selected for further use. The primer pairs tested are as follows: *Zrf1* (PP12004: 81.2%, PP24706: 80.3%, PP36474: 96.0%), *RpL32* (PD41810: 99.6%), *myc* (PP17271: 75.9%, PP29594:

102.1%, PP4159: 94.9%). Primers for rtPCR are mostly derived from *Hong et al. 2016* and are as follows:^31^

Stellate-F: GGCGATGAAAAGAAGTGGTA

Stellate-R: GCAGCGAGAAGAAGATGTCC

springer-F: CCATAACACCAGGGGCA

springer-R: CGAGTGCTGGTCTGTCA

ninja-F: GCCATCACCGACTACCATTA

ninja-R: TCACCCCATCCGACGCTCTT

roo-F: CGTCTGCAATGTACTGGCTCT

roo-R: CGGCACTCCACTAACTTCTCC

I-element-F: CAATCACAACAACAAAATC

I-element-R: GGTGTTGGTGTGGTTGGTTG

Pogo-F: CCAAAAATTTAACGACGCCTTT

Pogo-R: GCGCCAGCGCCAAA

hobo-F: CAAGTGCGACCGTCGACAT

hobo-R: AAGTGATGCCCAAAAAGTTTCTTT

copia-F: GAGGTTGTGCCTCCACTTAAGATT

copia-R: CAATACCACGCTTAGTGGCATAAA

NOF-F: TGCATCGAAGCTGTTTGCA

NOF-R: GAGTTTTAACGGTCTTGCGTTTC

R1-F: TGCGCCTACGGTGAGGTT

R1-R: CCGACAGCTACGCACATACG

blood-F: TATCGCATGGCAGATAGCCAAA

blood-R: CGTGGAATTCGGAAGTGGTTTC

frogger-F: GAAATACAGATCCTCACCACGAAGA

frogger-R: AGGAATGCATATCCTGAATACGACT

accord-F: TTTCATTCAAACAATTCCCTTACCT

accord-R: AGTTATGTAGTCCATTTTCCCGGTT

R2-F: GGGTGACCCTTTGTCTCCTATTCTA

R2-R: CAACGTCTTGTCCAACAATACTTGA

412-F: AAAGTACGGTCCAATGAAGACG

412-R: GTGGTGATGAGCTGTTGATGTT

Mdg1-F: CTTCAGTACCCAGATTTCAGCAAA

Mdg1-R: CCGCTCCACATGCTTGTTTA

G3-F: ACCATACCAGTTAGTGAAACGGTGA

G3-R: CTGCTTTATACAGCCGTAATTTGTTC

297-F: CTGGCAAAGGGATTTCATCA

297-R: TGCATTCCTAAGGCCAAATG

Gypsy-F: CAACAATCTGAACCCACCAATCT

Gypsy-R: TATGAACATCATGAGGGTGAACG

ZAM-F: ACTTGACCTGGATACACTCACAAC

ZAM-R: GAGTATTACGGCGACTAGGGATAC

IVK-F: ATATCAAAACCTAACAAACCCCCTT

IVK-R: CGTCCATCGTGCTATGATTTCTT

Blastopia-F: TTGTAGCTGGCCGCAAGAT

Blastopia-R: TTGACCCCTGTGACTTCGTATG

1731-F: TATGGGCTGAGGCGATAAAC

1731-R: CAAGTGGCTCACTGCTGGTA

Het-A-F: CGCGCGGAACCCATCTTCAGA

Het-A-R: CGCCGCAGTCGTTTGGTGAGT

I-element-F: CAATCACAACAACAAAATC

I-element-R: GGTGTTGGTGTGGTTGGTTG

TART-F: GATAATAGCCGATAAGCCCGCCA

TART-R: AAGACACAGCGGTTGATCGATATG

Cdk7-F: GGGTCAGTTTGCCACAGTTT

Cdk7-R: GATCACCTCCAGATCCGTG

### Cell Culture and RNA interference

*Drosophila* S2R+ cells were cultured at 25°C in Schneider’s medium with 10% fetal bovine serum (FBS) and streptomycin. HEK293T cells were cultured at 37°C in DMEM (Invitrogen 10-017-CV) supplemented with 10% FBS and streptomycin. For RNAi experiments, DNA templates for dsRNA were obtained from the *Drosophila* RNAi screening center (DRSC). Amplicons used in this study were: *Zrf1*(#11620), *Myc* (#23360, #37533), *Fmr1* (#26781), *vig* (#03631), *vig2* (#14402), *AGO2* (#10847), *Rm62* (#25109), *Tdrd3* (#10016) and *LacZ* dsRNA. A second dsRNA *Zrf1* template was generated by amplifying *Zrf1* cDNA with primers containing overhanging T7 sequences (JL278 and JL279). DNA templates were amplified with T7 primers using Phusion polymerase. PCR products were *in vitro* transcribed (IVT) using MEGAscript T7 Transcription Kit (ThermoFisher, AMB1334-5). S2R+ were treated with dsRNAs by the “bathing” method (https://fgr.hms.harvard.edu/drsc-cellrnai).

JL278: TAATACGACTCACTATAGGGTCTGGAGCAATGAGAACGTG

JL279: TAATACGACTCACTATAGGGGGTTGACTGATTGTTCGCCT

### Cell viability

1x10^5^ cells were plated into 96-well plates. Treated with 500ng of dsRNAs in a total volume of 80ul. 10ul was used and counted for cell number with a haemocytometer.

### Plasmids. Molecular cloning – *Drosophila* constructs

*<u>pEntr-GFP-Stop</u>*: The *pEntr-dTOPO* backbone was amplified from *pEntr-Akt* (Gift from P. Saavedra) with the primers pEntr_F & pEntr_R. GFP was amplified from the plasmid *pAW-vhh05-GFP* (gift from J. Xu) with JL201 and JL202. The two fragments were joined by Gibson assembly (NEB, E2611S). *<u>pEntr-GFP-Zrf1-Stop:</u>* The backbone of *pEntr-GFP-stop* was PCR amplified with the primers pEntr_F & JL203. The *Zrf1* coding sequence was amplified from the vector LD23875 with JL204 & JL205. The linker (GGGSGSG) was introduced between the GFP and *Zrf1* by overhanging sequences on the primers. Amplicons were joined by Gibson assembly. *<u>pEntr-Zrf1-NoStop:</u>* The *pEntr-dTOPO* backbone was amplified from *pEntr-GFP-stop* with the pEntr_F & pEntr_R. The *Zrf1*coding sequence was amplified from the plasmid LD23875 with JL171 & JL172 (primers that omitted the stop codon). Amplicons were joined by Gibson assembly. *<u>pEntr-AGO2-RB-NoStop</u>*: The AGO2 coding sequence was amplified from the plasmid *pAFW-AGO2* (Addgene #50554) with the primers JL267 & JL268. The fragment was inserted into the *pEntr* backbone via Gibson assembly. *<u>pEntr-FDY-NoStop</u>*: Two gBlock (IDT) dsDNA fragments (gJL1 & gJL2) encoding the entire *FDY* coding region (Accession KR781487) were joined with the *pEntr* backbone using Gibson assembly. *<u>pEntr-Tdrd3-NoStop</u>*: The *Tdrd3* coding sequence was amplified from the plasmid RE1471 with JL293 & 294. *<u>Final vectors:</u> pEntr223* vectors containing full-length cDNAs of *dMyc*, *Hspa14 (CG7182)*, *Zc3h15* (*CG8635*), *Hsc70-4*, *mod*, *peng*, *Fkbp39*, *Rm62*, *rin*, *Fmr1*, *vig*, *vig2, Ref1, RpS3, 128up, CG8584* and *yps* were from the FlyBi ORFeome Collection. The *pEntr-mCherry-NoStop* was a gift from B. Xia. All fly *pEntr*vectors containing cDNAs were transferred into the *Drosophila* gateway vectors *pAW, pAWH, pAWM* and *pAGW* (Carnegie Science/Murphy lab) using LR clonase II enzyme mix (Invitrogen, 11791-020) (see table for full list of vectors).

JL171: CCGCGGCCGCCCCCTTCACCATGACGAGCGGTACGGTAGC

JL172: GGGTCGGCGCGCCCACCCTTTTTGACCGCCGCCTGTG

JL201: CCGCGGCCGCCCCCTTCACCATGGTGAGCAAGGGCGAG

JL202: GGGTCGGCGCGCCCACCCTTTCACTTGTACAGCTCGTCCATGCC

JL203: GCCGGAGCCGCTGCCACCTCCCTTGTACAGCTCGTCCATGC

JL204: GGAGGTGGCAGCGGCTCCGGCATGACGAGCGGTACGGTAGC

JL205: GGGTCGGCGCGCCCACCCTTTCATTTGACCGCCGCCTG

JL267: CCGCGGCCGCCCCCTTCACCATGGGAAAAAAAGATAAGAACAAGAAAGGAG

JL268: GGGTCGGCGCGCCCACCCTTGACAAAGTACATGGGGTTTTTCTTCA

JL293: CCGCGGCCGCCCCCTTCACCATGGAATTAGGCAAGAAACTACG

JL294: GGGTCGGCGCGCCCACCCTTATGCTCTCGTTTGTGGG

### Molecular cloning – Human constructs

*<u>pEntr-ZRF1-NoStop</u>*: The human *ZRF1* coding sequence was amplified from the plasmid *pcDNA3.1-ZRF1* (GenScript, OHu13274) with JL186 & JL188. *<u>pEntr-SERBP1-NoStop:</u>* The human SERBP1 coding sequence was amplified from the plasmid UAS-SERBP1 (DGRC, hGUHO03824) with JL313 & JL314. *<u>Final vectors:</u> pDonr221-HABP4-NoStop* (DNASU HsCD00745639), *pEntr-mCherry-NoStop* and other *pEntr* vectors containing cDNAs were transferred into the mammalian gateway vectors, *pHAGE-N-FLAG-HA* (*pHFH;* gift from R. Viswanatha) or pCSF107mT-GATEWAY-3’LAPtag (*pCSF-GFP*, Addgene #67618, gift from S. Entwisle) using LR clonase II enzyme mix. Other plasmids containing epitope-tagged cDNAs were used and are listed as follows*: pDEST-myc-DDX17* (Addgene #19876).

JL186: CCGCGGCCGCCCCCTTCACCATGCTGCTTCTGCCAAGCG

JL188: GGGTCGGCGCGCCCACCCTTTCATTTCTTGGCTCTACTTGCATTCAGCA

JL313: CGCGGCCGCCCCCTTCACCATGCCTGGGCACTTACAGGAAG

JL314: GGGTCGGCGCGCCCACCCTTAGCCAGAGCTGGGAATGCC

### Immunoprecipitation and immunoblotting

Immunoprecipitation was performed as previously described.^40^ S2R+ or HEK293T cells were split 2 days before transfection. 2ug of DNA was transfected into 20 x 10^6^ of cells in a 100mm dish with Effectene transfection reagent (Qiagen 301427) following the manufacturer’s protocol. After 3 days, cells were lysed with IP-lysis buffer (Pierce 87787) containing 2X protease inhibitor cocktail (ThermoFisher, 78430) and 2X PhosSTOP (Sigma-Aldrich 4906837001). Lysate was incubated with 25ul of Chromotek-GFP-trap beads (Bulldog Biotechnology gta-20) or anti-Flag agarose (Sigma A2220) and subject to end-over-end agitation for 2hrs at 4°Cs. Beads were washed 3-4 times with 500ul of lysis buffer. Beads were eluted in 25ul of sample buffer (Pierce 39001 with 10% 2-Mercaptoethanol), boiled for 5min and sedimented by centrifugation at 2500x g for 5min.

For immunoblotting, samples were run on 4%-20% polyacrylamide gel (Bio-Rad 4561096) and transferred to an Immobilon-P polyvinylidene fluoride (PVDF) 0.45um membrane (Millipore IPVH00010). The membrane was blocked with 5% skim milk in TBST (TBS with 0.05% Tween-20) at room temperature for 1h and probed with primary antibody in 5% skim milk overnight at 4°C. Membrane was probed with HRP-conjugated secondary antibody at room temperature for 1h and signal detected by enhanced chemiluminescence (ECL;Amersham RPN2209; Pierce 34095).

Concentrations used for primary antibodies were as follows: chicken anti-GFP (1:10000, aveslabs GFP-1020), use anti-HA.11 (1:1000, BioLegend 901514), mouse anti-Fmr1 (1:100, DSHB 5A11), rabbit anti-vig (1:30000, gift from N. Riddle) & rabbit-vig2 (1:10000, gift from N. Riddle), mouse anti-dAGO2 (1:400, gift from H. Siomi), rat anti-Rm62 (1:2000, gift from E. Lei), rabbit anti-Tdrd3 (1:2000, gift from W. Wang), mouse anti-lost (1:100, DSHB), mouse anti-Tubulin (1:1000, Sigma T5168).

### Measuring eye pigmentation

*w^m4h^, y^1^, ac*^1^ females were crossed with different alleles and reared at 25°C. In the progeny, the left eye of 2-day-old adult male flies were imaged and used for measuring eye pigmentation. Images were separated into the RBG channels, and the mean grey intensity of the eye was measured under the red channel. This value was subtracted from the background mean grey intensity and compared between genotypes as a fold-change of the *w^m4h^, y^1^, ac*^1^ control.

### Generating fly mutants with CRISPR

*<u>vig</u>* <u>mutants:</u> *y; P{TKO.GS04822}*attP40 (BL81492) was crossed with *y,w; nos-Cas9/CyO*. *y,w; nos-Cas9/ P{TKO.GS04822}*attP40 males were crossed with *y,w; Gla/CyO* females. Resulting offsprings were outcrossed individually to establish stocks. A ∼500bp region covering the seed sequence was amplified with JL317 & JL318 and subcloned into pJET1.2/blunt vector for sequencing. The selected mutants, *vig^d^*^65–72^ and *vig^d^*^63–73^ have predicted premature stop codons that result in respectively truncated proteins of 59aa and 58aa.

JL317: ATCGAATCTCACAGGGTGTGAGAG

JL318: GTTTGTTGTCGGCCGATGGA

*<u>vig2</u>* <u>mutants:</u> *y; P{TKO.GS04324}attP40* (BL84243) was crossed with *y,w; nos-Cas9/CyO*. *y,w; nos-Cas9/ P{TKO.GS04324}attP40* males were crossed with *y,w; TM3/TM6b* females. Resulting offsprings were outcrossed individually to establish stocks. A ∼600bp region covering the seed sequence was amplified with JL315 & JL316 and subcloned into pJET1.2/blunt vector for sequencing. The selected mutants, *vig2^d^*^53–72^, *vig2^d^*^57–66^ and *vig2^G.d^*^59–75^, have predicted premature stop codons that result in respectively truncated proteins of 61aa, 26aa and 25aa.

JL315: GCCATTGGGACCAAAGTAACTCTCAAA

JL316: TTACAGGGCAGCATTTTCCACG

### Single-nuclei bioinformatics analysis

Reads were mapped to the *Drosophila melanogaster* reference genome (BDGP6.46.111) using STARsolo (STAR version 2.7.9a). SoloTE (release 1.09) with default parameters was subsequently run to count and assign TE reads at the subfamily and loci-specific levels. Count matrices were then processed in R (version 4.3.2) using Seurat (version 5.0.2). Droplets which were identified as empty by CellRanger (version 7.1.0) were filtered out. Low quality nuclei were filtered out using the following parameters: mitochondrial percentage > 0.05%, number of genes < 300, Log10(number of genes/number of UMIs) < 0.80. The data was normalized using the NormalizeData function and 2,000 highly variable features were identified using the FindVariableFeatures function. The data was scaled, and dimensionality reduction was performed using principal component analysis (PCA) on the highly variable features. Using a graph-based clustering approach, the FindNeighbours and FindClusters functions were used (dims = 30) to identify clusters within the data.

Non-linear dimensional reduction was performed with UMAP for visualization of clusters. Using resolution 0.3, two clusters were split according to their behavior at higher resolution (0.8). The FindAllMarkers function was used to identify marker genes for cell type annotation of clusters. PCA was performed on gene only and TE only expression features for each cluster to explore the highest sources of variation within the data. Using a Wilcoxon rank-sum test, differentially expressed genes (DEGs) between conditions for each cell type were identified (abs(log2fc) > 0.38 & p-value < 0.05). These DEGs were then visualized on a volcano plot using ggplot2 (version 3.5.0). Gene set enrichment analysis using GO:BP terms was performed using the following ranking metric for each cell type: sign(log2fc) * -log10(p-value). Pathways with Benjamini-Hochberg adjusted p-values less than 0.05 were considered significantly enriched.

## Author contribution

JSSL conceived, designed and performed all *in vivo* and biochemical experiments. JSSL and JX collected virgins and male flies in the morning and afternoon, respectively. YL conducted all single nuclei related experiments with help from JSSL. JMA conducted and obtained the mass spectrometry data. YH analyzed mass spectrometry data. YH and MQ analyzed snRNA-seq data. BX generated the pEntr-mcherry vector and S2R+ cells stably expressing AWH-empty-HSP-GFP-puromycin. AK conducted the *in silico* interaction predictions with AFM. RB generated all competent cells used in this study and assisted in the husbandry of CRISPR mutant flies following crosses. NP supervised the project. JSSL wrote the paper with input from NP, AK and YH.

## Acknowledgements

We thank the Siomi lab, Nicole Riddle, the ModEncode project, Elissa Lei and Weidong Wang for antibodies. We thank Yassi Hafezi from the Clark Lab for providing FDY mutants. We thank Hyuckjoon Kang from the Kuroda lab for providing *Pc* mutants. We thank Paula Montero Llopis for providing training and technical assistance with confocal imaging at the Microscopy Resources on the North Quad (MicRoN) core at Harvard Medical School. We thank Perrimon lab members and Danesh Moazed for discussion and feedback. We thank Cathryn (King) Murphy, Litz Brown and Rich Binari for ordering and coordinating reagents for all experiments.

We thank Litz Brown editing and adding reference. JSSL was supported by a Postdoctoral Croucher fellowship from the Croucher Foundation of Hong Kong. NP is an investigator of Howard Hughes Medical institute. This article is subject to HHMI’s Open Access to Publications policy. HHMI lab heads have previously granted a non-exclusive CC BY4.0 license to the public and a sublicensable license to HHMI in their research articles. Pursuant to those licenses, the author-accepted manuscript of this article can be made freely available under a CC BY 4.0 license immediately upon publication.

---

**Figure 1:**

(B, C and J)

attP^VIE-260B^: *w*; *esg-Gal4, UAS-GFP, tub-Gal80^ts^/VIE-260b (KK control); +/+*

attP40: *w*; *esg-Gal4, UAS-GFP, tub-Gal80^ts^/attP40 (empty); +/+*

Luc-i: *w*; *esg-Gal4, UAS-GFP, tub-Gal80^ts^/+; UAS-Luc-i (JF01355)/+*

Zrf1-i #1: *w*; *esg-Gal4, UAS-GFP, tub-Gal80^ts^/UAS-Zrf1-i (HMC05212); +/+* Zrf1-i #2: *w*; *esg-Gal4, UAS-GFP, tub-Gal80^ts^/UAS-Zrf1-i (GL01550); +/+* Zrf1-i #3: *w*; *esg-Gal4, UAS-GFP, tub-Gal80^ts^/+; UAS-Zrf1-i (10565R-1)/+* Zrf1-i #4: *w*; *esg-Gal4, UAS-GFP, tub-Gal80^ts^/UAS-Zrf1-i (KK102408); +/+* Zrf1-i #5: *w*; *esg-Gal4, UAS-GFP, tub-Gal80^ts^/+; UAS-Zrf1-i (GD7019)/+* p35: *w*; *esg-Gal4, UAS-GFP, tub-Gal80^ts^/+; UAS-p35/+*

Zrf1-i #1, p35: *w*; *esg-Gal4, UAS-GFP, tub-Gal80^ts^/UAS-Zrf1-i (HMC05212); UAS-p35/+*

DIAP: *w*; *esg-Gal4, UAS-GFP, tub-Gal80^ts^/+; UAS-DIAP/+*

Zrf1-i #1, DIAP: *w*; *esg-Gal4, UAS-GFP, tub-Gal80^ts^/UAS-Zrf1-i (HMC05212); UAS-DIAP/+*

(D and K) *w*; *esg-Gal4, UAS-GFP, tub-Gal80^ts^/+; UAS-Luc-i (JF01355)/Dl-LacZ* (E and K) *w*; *esg-Gal4, UAS-GFP, tub-Gal80^ts^/UAS-Zrf1-i (HMC05212); Dl-LacZ/+* (F and J) *w*; *esg-Gal4, UAS-GFP, tub-Gal80^ts^/+; UAS-FLP, act>CD2>Gal4/UAS-Luc-i (JF01355)*

(G and J) *w*; *esg-Gal4, UAS-GFP, tub-Gal80^ts^/UAS-Zrf1-i (HMC05212); UAS-FLP,*

*act>CD2>Gal4/+*

(H and J) *w*; *esg-Gal4, UAS-GFP, tub-Gal80^ts^/UAS-sgZrf1 (CFD00331); UAS-Cas9.P2/+*

(I and J) *w*; *esg-Gal4, UAS-GFP, tub-Gal80^ts^/UAS-sgIntergenic (BEC); UAS-Cas9.P2/+*

(L) [from left to right]^30^

*w*; *esg-Gal4, UAS-GFP, tub-Gal80^ts^, Su(H)-Gal80/+; UAS-Luc-i (JF01355)/+*

*w*; *esg-Gal4, UAS-GFP, tub-Gal80^ts^, Su(H)-Gal80/UAS-Zrf1-i (HMC05212); +/+*

*w*; *Su(H)GBE-Gal4/+; tub-Gal80^ts^/UAS-Luc-i (JF01355)*

*w*; *Su(H)GBE-Gal4/UAS-Zrf1-i (HMC05212); tub-Gal80^ts^/+ w*; *tub-Gal80^ts^/+; pros-Gal4/UAS-Luc-i (JF01355)*

*w*; *tub-Gal80^ts^/UAS-Zrf1-i (HMC05212); pros-Gal4/+*

**Figure 2:**

(A and B)

w^1118^: *w*; *esg-Gal4, UAS-GFP, tub-Gal80^ts^/+; +/+* (*egt^ts^* crossed with *w^1118^*) Zrf1^P{EP}G4964^: *w*; *esg-Gal4, UAS-GFP, tub-Gal80^ts^/+; Zrf1^P{EP}G4964^/+*

Zrf1^P{EPgy2}G4964^: *w*; *esg-Gal4, UAS-GFP, tub-Gal80^ts^/+; Zrf1^P{EPgy2}G4964^/+* Zrf1^ORF.3xHA^: *w*; *esg-Gal4, UAS-GFP, tub-Gal80^ts^/+; Zrf1^ORF.3xHA^/+* Zrf1^ORF.CC^: *w*; *esg-Gal4, UAS-GFP, tub-Gal80^ts^/+; Zrf1^ORF.CC^/+*

(C and D)

attp40: *w*; *esg-Gal4, UAS-GFP, tub-Gal80^ts^/attP40 (empty); UAS-dCas9-VPR/+*

sg-Zrf1: *w*; *esg-Gal4, UAS-GFP, tub-Gal80^ts^/sg-Zrf1(GS03041); UAS-dCas9-VPR/+*

(E-I)

w^1118^: *w*; *esg-Gal4, UAS-GFP, tub-Gal80^ts^/+; +/+* (*egt^ts^* crossed with *w^1118^*) Egfr^Top4.2^: *w/UAS-Egfr^Top4.2^*; *esg-Gal4, UAS-GFP, tub-Gal80^ts^/+; +/+* Egfr^Top4.2^+Zrf1-i: *w/UAS-Egfr^Top4.2^*; *esg-Gal4, UAS-GFP, tub-Gal80^ts^/UAS-Zrf1-i (HMC05212); +/+*

Ras1^A^: *w*; *esg-Gal4, UAS-GFP, tub-Gal80^ts^/+; UAS-Ras1^A^/+*

Ras1^A^+Zrf1-i: *w*; *esg-Gal4, UAS-GFP, tub-Gal80^ts^/UAS-Zrf1-i (HMC05212); UAS-Ras1^A^/+*

Raf^F179^: *w*; *esg-Gal4, UAS-GFP, tub-Gal80^ts^/+; UAS-Raf^F179^/+*

Raf^F179^+Zrf1-i: *w*; *esg-Gal4, UAS-GFP, tub-Gal80^ts^/UAS-Zrf1-i (HMC05212); UAS-Raf^F179^/+*

hRaf^gof^: *w*; *esg-Gal4, UAS-GFP, tub-Gal80^ts^/+; UAS-hRaf^gof^/+*

hRaf^gof^+Zrf1-i: *w*; *esg-Gal4, UAS-GFP, tub-Gal80^ts^/+; UAS-hRaf^gof^/+*

(J)

Luc-i^JF01355^: *w*; *esg-Gal4, UAS-GFP, tub-Gal80^ts^/+; UAS-Luc-i (JF01355)/+*

N-i^HMS00009^: *w*; *esg-Gal4, UAS-GFP, tub-Gal80^ts^/+; UAS-N-i (HMS00009)/+*

N-i^HMS00009^+ Zrf-i: *w*; *esg-Gal4, UAS-GFP, tub-Gal80^ts^/UAS-Zrf1-i (HMC05212); UAS-N-i (HMS00009)/+*

N-i^JF01637^: *w*; *esg-Gal4, UAS-GFP, tub-Gal80^ts^/+; UAS-N-i (JF01637)/+*

N-i^JF01637^+Zrf1-i: *w*; *esg-Gal4, UAS-GFP, tub-Gal80^ts^/UAS-Zrf1-i (HMC05212); UAS-N-i (JF01637)/+*

N-i^HMS00001^: *w*; *esg-Gal4, UAS-GFP, tub-Gal80^ts^/+; UAS-N-i (HMS00001)/+*

N-i^HMS00001^+Zrf1-i: *w*; *esg-Gal4, UAS-GFP, tub-Gal80^ts^/UAS-Zrf1-i (HMC05212); UAS-Luc-i (HMS00001)/+*

(K)

w^1118^: *w*; *esg-Gal4, UAS-GFP, tub-Gal80^ts^/+; +/+* (*egt^ts^* crossed with *w^1118^*) yki^WT^: *w*; *esg-Gal4, UAS-GFP, tub-Gal80^ts^/+; UAS-yki^WT^/+*

yki^WT^+Zrf1-i: *w*; *esg-Gal4, UAS-GFP, tub-Gal80^ts^/UAS-Zrf1-i (HMC05212); UAS-yki^WT^/+*

yki^3SA^: *w*; *esg-Gal4, UAS-GFP, tub-Gal80^ts^/+; UAS-yki^3SA^/+*

yki^3SA^+Zrf1: *w*; *esg-Gal4, UAS-GFP, tub-Gal80^ts^/UAS-Zrf1-i (HMC05212); UAS-yki^3SA^/+*

**Figure 4:**

Genotypes are as listed in the figure.

**Figure 5:**

(C and D) y,w/Y: *y,w/Y* OreR/Y: *OreR/Y*

Control: *w^m4h^,ac^1^,y^1^/Y* (*w^m4h^,ac^1^,y*^1^ females crossed with *y,w* males) Fmr1^d50M^/+: *w^m4h^,ac^1^,y^1^/Y; +/+; Fmr1^d50M^/+*

Fmr1^d113M^/+: *w^m4h^,ac^1^,y^1^/Y: +/+ : Fmr1^d113M^/+*

vig^C274^/+: *w^m4h^,ac^1^,y^1^/Y; vig^C274^/+; +/+* vig^d65–72^/+: *w^m4h^,ac^1^,y^1^/Y;* vig^d65–72^/+; +/+ vig^d63–73^/+: *w^m4h^,ac^1^,y^1^/Y; vig^d^*^63–73^*/+; +/+*

vig2^PL00470^/+: *w^m4h^,ac^1^,y^1^/Y; +/+; vig2^PL00470^/+*

vig2^Gd59–75^/+: *w^m4h^,ac^1^,y^1^/Y; +/+; vig2^Gd^*^59–75^*/+*

FDY^WT_E5^: *w^m4h^,ac^1^,y^1^/Y, FDY^WT_E5^* (control for FDY mutants)

FDY^MUT_L3^: *w^m4h^,ac^1^,y1/Y, FDY^MUT_L3^*

FDY^MUT_L64^: *w^m4h^,ac^1^,y^1^/Y, FDY^MUT_L64^*

AGO2^414^/+: *w^m4h^,ac^1^,y^1^/Y; +/+; AGO2^414^/+*

Rm62^01086^/+: *w^m4h^,ac^1^,y^1^/Y: +/+; Rm62^01086^/+*

BSC553/+: *w^m4h^,ac^1^,y^1^/Y: +/+; BSC553/+*

Zrf1^G4964^/+: *w^m4h^,ac^1^,y^1^/Y: +/+; Zrf1^G4964^/+*

Zrf1^cr01803^/+: *w^m4h^,ac^1^,y^1^/Y: +/+; Zrf1^cr01803^/+*

Zrf1^016^/+: *w^m4h^,ac^1^,y^1^/Y: +/+; Zrf1^016^/+*

Zrf1^018^/+: *w^m4h^,ac^1^,y^1^/Y: +/+; Zrf1^018^/+*

Zrf1^127^/+: *w^m4h^,ac^1^,y^1^/Y: +/+; Zrf1^127^/+*

Zrf1^148^/+: *w^m4h^,ac^1^,y^1^/Y: +/+; Zrf1^148^/+*

Zrf1^242^/+: *w^m4h^,ac^1^,y^1^/Y: +/+; Zrf1^2^*^42^*/+*

(E, F and G)

Genotypes are as listed in the figure.

**Figures S1:**

(B, C and D)

Luc-i: *w*; *esg-Gal4, UAS-GFP, tub-Gal80^ts^/+; UAS-FLP, act>CD2>Gal4/UAS-Luc-i (JF01355)*

Zrf1-i: *w*; *esg-Gal4, UAS-GFP, tub-Gal80^ts^/UAS-Zrf1-i (HMC05212); UAS-FLP,*

*act>CD2>Gal4/+*

(E)

Luc-i: *w*; *tub-Gal80^ts^/+; Dl-Gal4/UAS-Luc-i (JF01355)*

Zrf1-i: *w*; *tub-Gal80^ts^/UAS-Zrf1-i (HMC05212); Dl-Gal4/+*

**Figure S6 and S7:**

Genotypes are as listed in the figure.

**Supplemental Figure 1:**
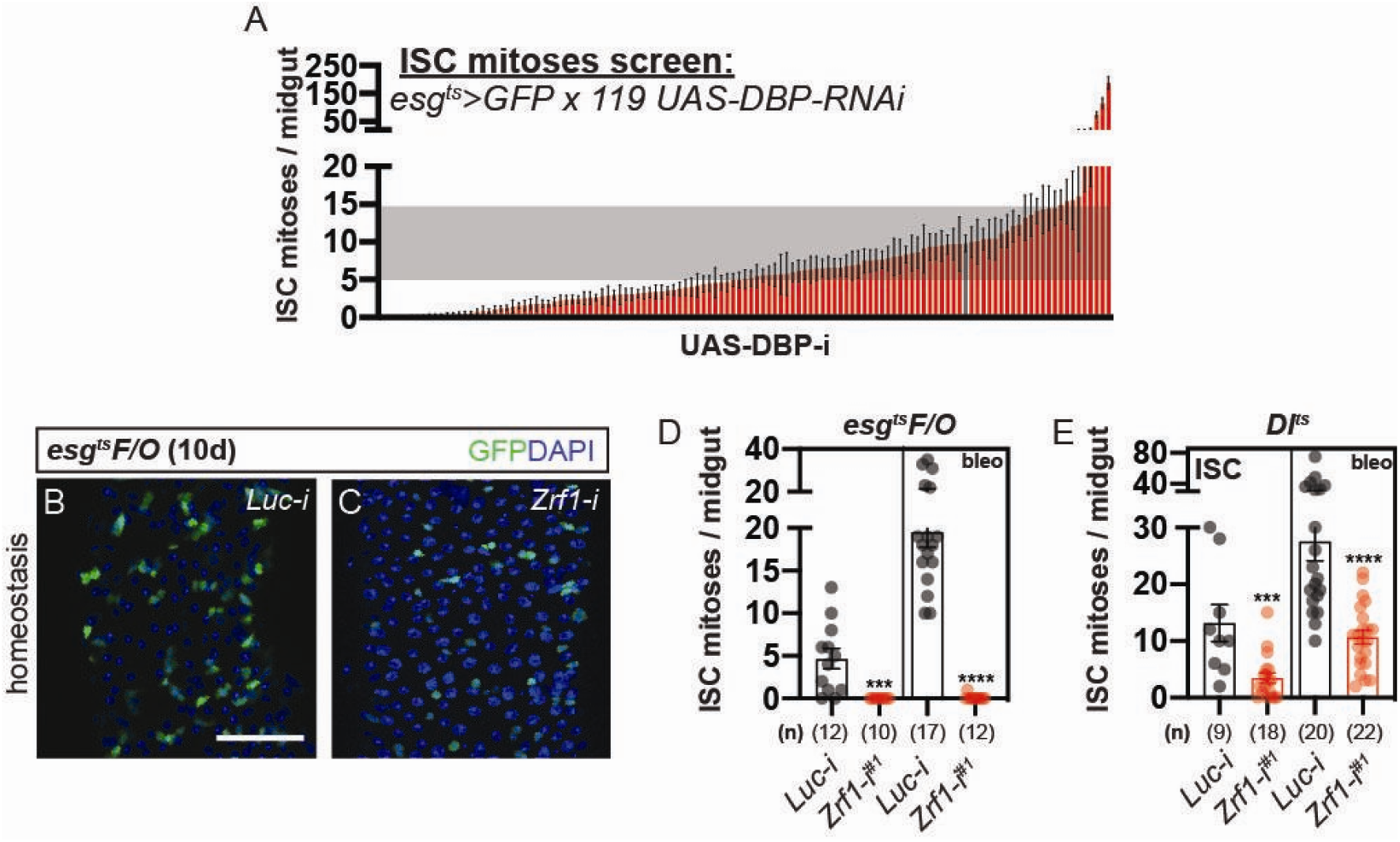
Midgut progenitor RNAi Screen Identifies *Zrf1*. (A) ISC mitoses per midgut from an RNAi screen targeting 119 DNA-binding proteins (*UAS-DBP-RNAi*) driven by *esg^ts^>GFP*. Each bar represents ISC mitoses for a specific RNAi line after 10 days at the permissive temperature. Red bars indicate lines with significantly reduced ISC mitoses compared to control (Luc-i). *Zrf1* RNAi (*Zrf1-i*) is among the top hits causing a strong reduction in ISC proliferation. (B–C) Confocal images showing GFP+ ISC/EB progenitor cells after 10 days of RNAi induction in *esg^ts^F/O* flies during homeostasis. (B) Control (*Luc-i*) showing normal ISC proliferation, and (C) *Zrf1* knockdown (*Zrf1-i*) showing a marked reduction in GFP+ progenitor cells. Green: GFP; blue: DAPI. Scale bar: 50 µm. (D) Quantification of ISC mitoses per midgut from *esg^ts^F/O* flies after 10 days of RNAi induction during homeostasis or bleomycin (bleo) treatment. (E) ISC mitoses per midgut from *DI^ts^* flies where *Zrf1* RNAi was specifically driven in ISCs. ISC proliferation is markedly reduced upon *Zrf1* knockdown compared to control (*Luc-i*), both during homeostasis and after bleomycin treatment. Bars represent mean ± SEM; n = number of midguts analyzed. Statistical significance determined by Student’s t-test; ***p < 0.001, ****p < 0.0001.*

**Supplemental Figure 2:**
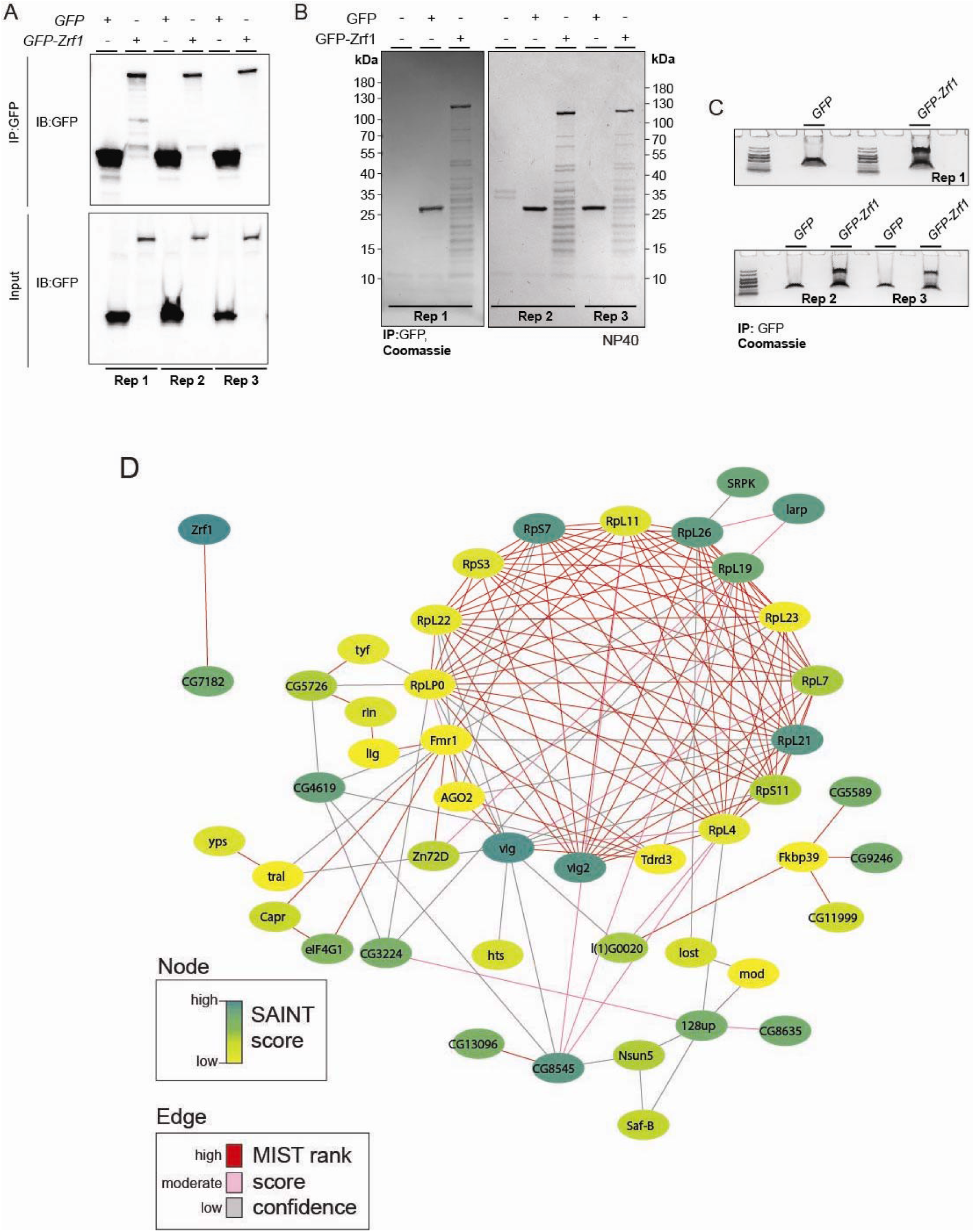
Validation of Zrf1 Interaction Network by IP-MS and Network Analysis. (A) Western blot analysis of immunoprecipitated (IP) samples from S2R+ cells expressing GFP or GFP-Zrf1 across three biological replicates (Rep 1, Rep 2, Rep 3). IP was performed using GFP-trap beads, and the presence of GFP-tagged proteins was confirmed by immunoblotting (IB) for GFP. Input samples show consistent expression levels across all replicates. (B) Coomassie-stained gels of IP samples from S2R+ cells expressing GFP or GFP-Zrf1. Three biological replicates (Rep 1, Rep 2, Rep 3) are shown, demonstrating consistent patterns of protein bands enriched in the GFP-Zrf1 lanes, confirming the reproducibility of Zrf1-associated protein complexes. The samples were prepared under mild detergent (NP40) conditions. (C) Additional Coomassie-stained gels highlighting the distinct protein bands observed in GFP-Zrf1 IP samples across the three replicates (Rep 1, Rep 2, Rep 3). This suggests specific enrichment of proteins that associate with Zrf1. (D) Network diagram showing the Zrf1 PPI network derived from IP-MS data. Nodes represent proteins, with colors indicating the SAINT score (green: high, yellow: low). Edges represent interactions, with line colors indicating MIST rank confidence (red: high, pink: moderate, gray: low). The network reveals associations between Zrf1 and multiple ribosomal proteins, RNA-binding proteins, and components of the RNA-induced silencing complex (RISC), highlighting the diverse interactome of Zrf1. IP-MS data validation was performed across three independent biological replicates, confirming reproducible interaction profiles. The network was constructed using SAINT and MIST scoring to assess interaction confidence.

**Supplemental Figure 3:**
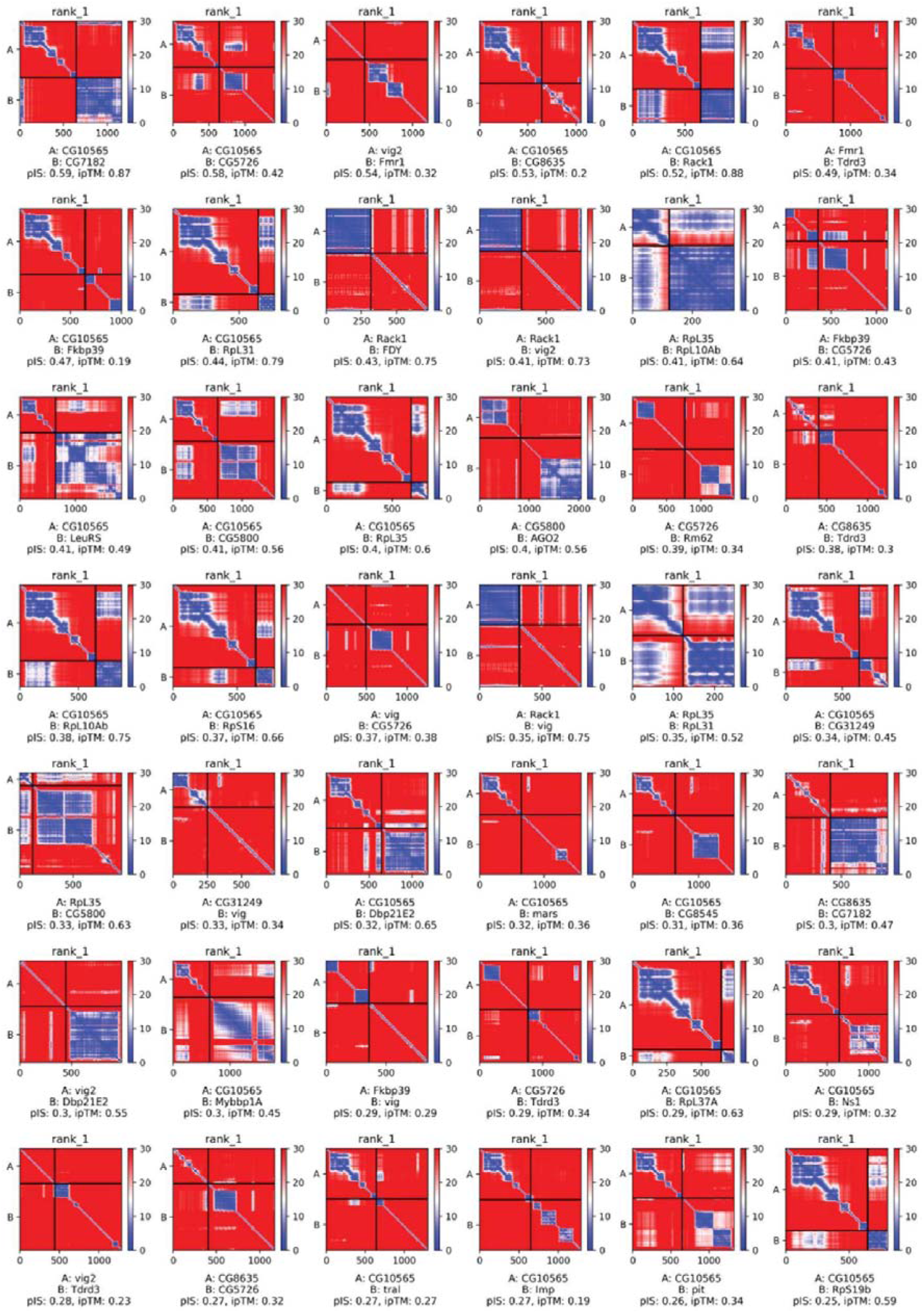
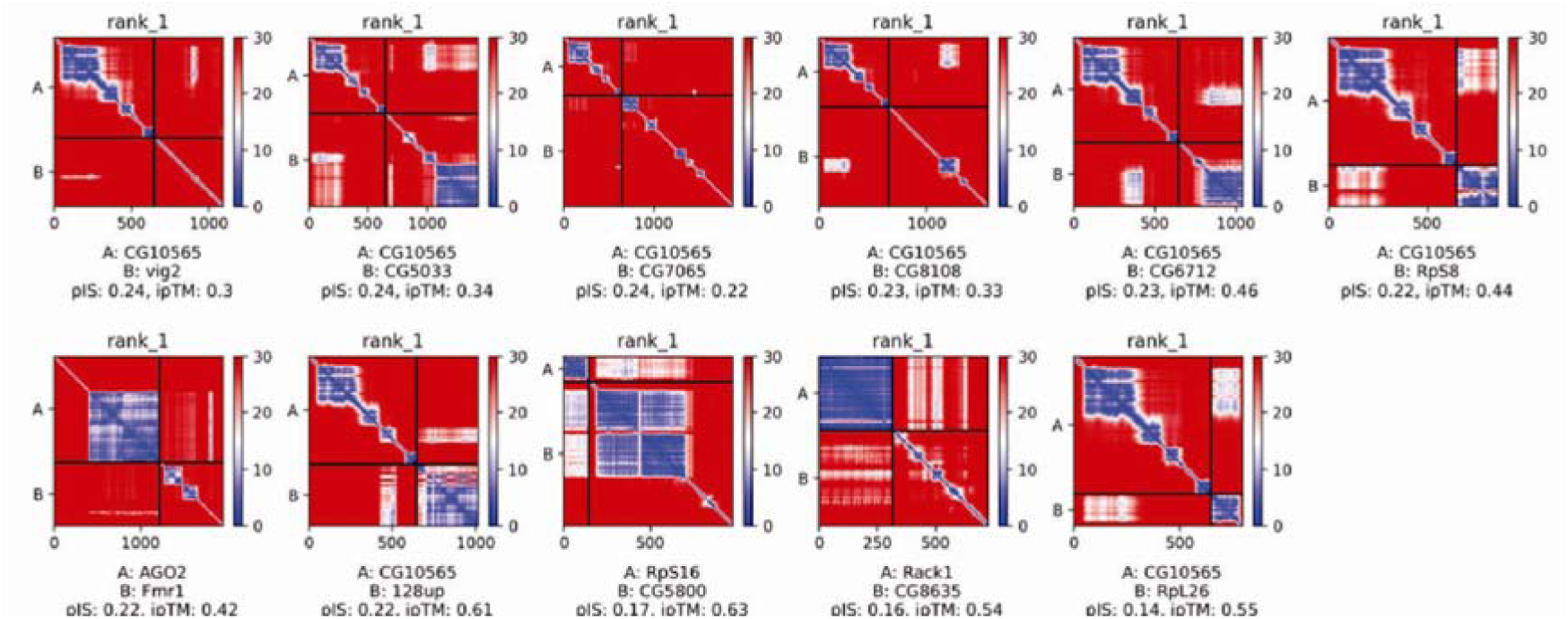
AlphaFold-Multimer predicted aligned error (PAE) maps for Zrf1 Interactome Predictions. PAE maps of positive PPIs using AlphaFold-Multimer (AFM) screens to predict direct interactions between Zrf1 (CG10565) and candidate interactors identified from IP-MS data. Each PAE map represents the residue-residue alignment confidence between Zrf1 and a candidate interactor. In these PAE maps, blue regions indicate low predicted alignment error, suggesting high confidence in residue-residue alignment (potential interaction), while red regions denote high alignment error, indicating lower confidence. Local Interaction Score (LIS), Local Interaction Area (LIA), and ipTM scores are displayed for each interaction. The cut-off criteria to classify positive PPIs were set at LIS ≥ 0.203 and LIA ≥ 3432. Some interactions identified by AFM predictions were further experimentally validated through Co-IPs. These AFM predictions provided a preliminary model for the Zrf1 interaction network and guided the subsequent experimental validation of Zrf1-associated proteins (Figure 3C).

**Supplemental Figure 4:**
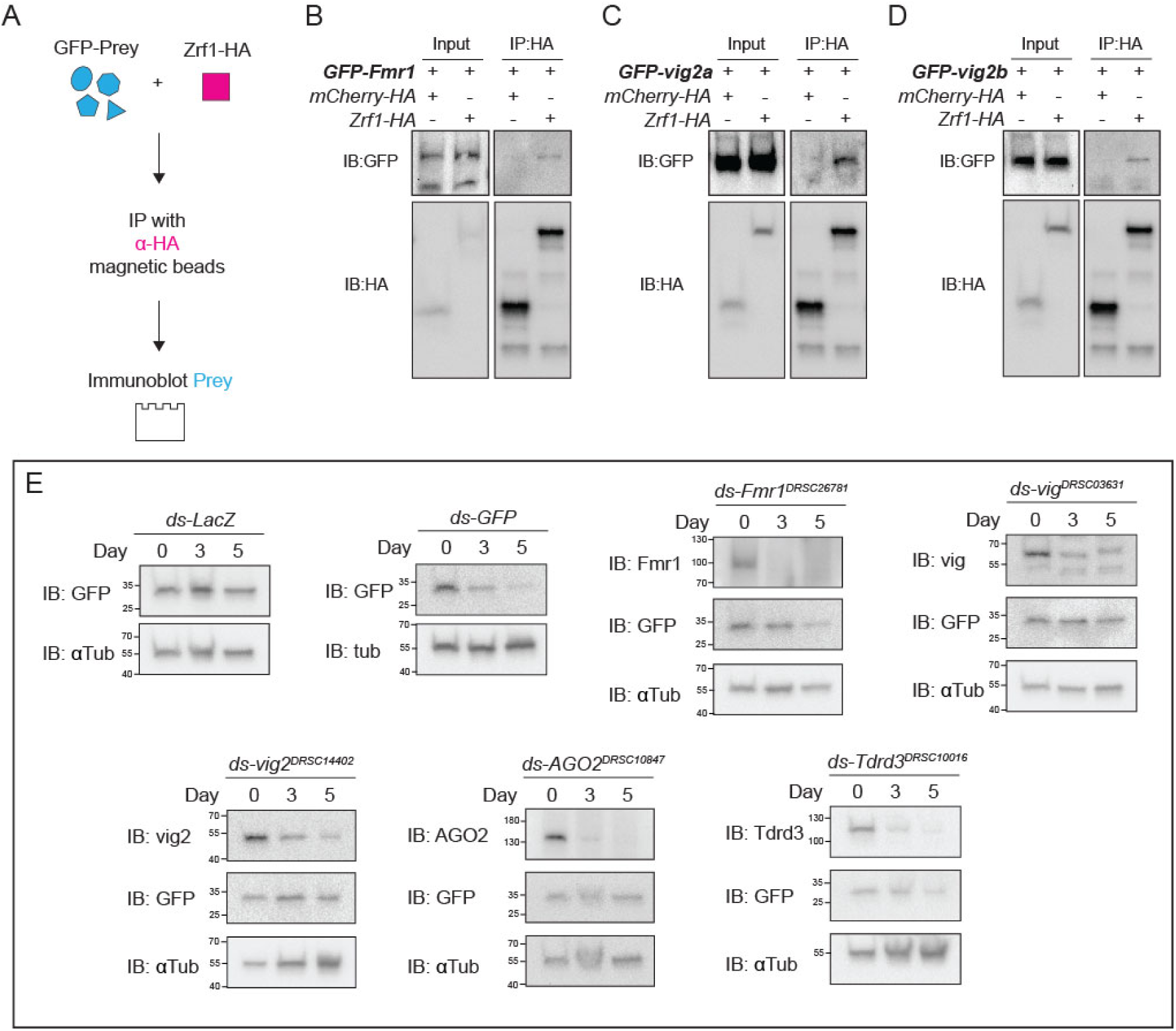
Validation of Zrf1 Interactions and Effects of RISC Component Knockdowns on GFP Expression. (A) Schematic of the co-immunoprecipitation (Co-IP) workflow. GFP-tagged prey proteins were co-expressed with Zrf1-HA, followed by immunoprecipitation using α-HA magnetic beads. Immunoblots were used to detect GFP-tagged prey proteins, verifying interactions. (B–D) Co-IP experiments confirming interactions between Zrf1-HA and GFP-tagged RISC components (B) Zrf1-HA interacts with GFP-Fmr1, showing co-precipitation after immunoprecipitation with α-HA beads. mCherry-HA was used as a negative control. (C) Interaction between Zrf1-HA and GFP-vig2a, with mCherry-HA as the control. (D) Co-IP showing that Zrf1-HA also interacts with GFP-vig2b. Inputs verify protein expression levels, and controls demonstrate specificity of the Co-IP. (E) Immunoblots showing the specificity of antibodies against RISC components. RNAi-mediated knockdown of various RISC components on GFP expression over time. S2R+ cells were treated with dsRNA against *LacZ* (control), *GFP, Fmr1, vig, vig2, AGO2,* and *Tdrd3* for 0, 3, and 5 days. Western blots display levels of target proteins (e.g., Fmr1, vig, vig2, AGO2, Tdrd3) and GFP. Knockdown efficiency is confirmed by the reduction of the respective proteins, while α-Tubulin (αTub) serves as a loading control.

**Supplemental Figure 5:**
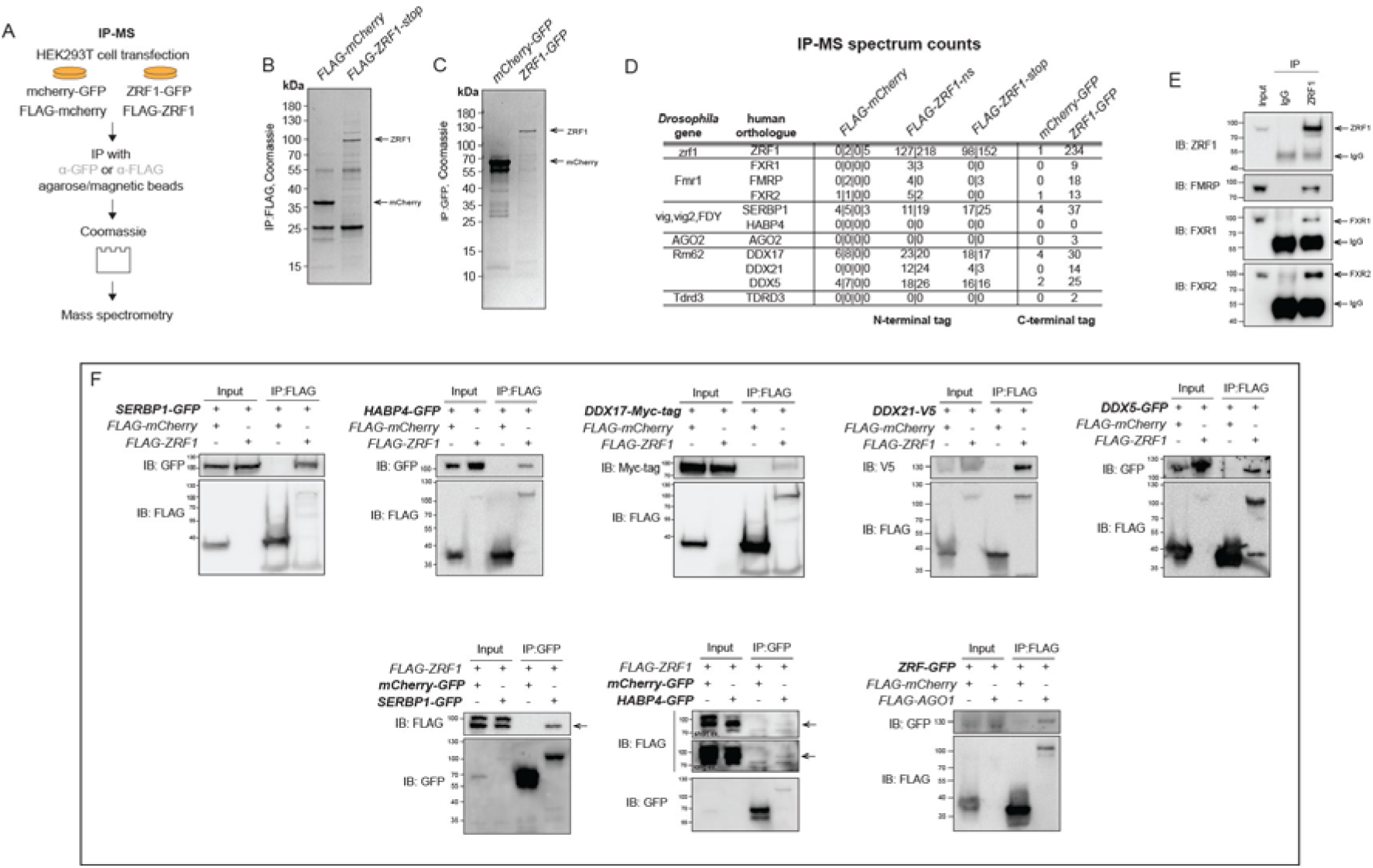
Protein-protein Interaction of Human ZRF1. (A) Schematic of the IP-MS workflow used to identify human ZRF1 interactors. HEK293T cells were transfected with mCherry-GFP, ZRF1-GFP, FLAG-mCherry, or FLAG-ZRF1. Immunoprecipitations (IPs) were performed using α-GFP or α-FLAG agarose/magnetic beads, followed by Coomassie staining and mass spectrometry analysis. (B) Coomassie-stained gel showing proteins immunoprecipitated using FLAG-tagged constructs (FLAG-mCherry, FLAG-ZRF1-stop). The FLAG-ZRF1-stop lane shows distinct protein bands, indicating specific enrichment of ZRF1-associated proteins. (C) Coomassie-stained gel displaying IP results for GFP-tagged constructs (mCherry-GFP, ZRF1-GFP). Bands in the ZRF1-GFP lane highlight the enrichment of ZRF1 and associated proteins. (D) Table summarizing the IP-MS spectral counts for human ZRF1 and known orthologs of *Drosophila* interactors. The table compares counts across different IP conditions (N-terminal vs. C-terminal tags) for FLAG-mCherry, FLAG-ZRF1, mCherry-GFP, and ZRF1-GFP, indicating consistency of interaction data across tagging strategies. Key interactors include FXR1, FXR2, DDX17, and SERBP1, which show orthology to *Drosophila* RISC components. (E) IPs with endogenous ZRF1 from HEK293T cells followed by immunoblotting for FMRP, FXR1, and FXR2, confirming interactions. Input lanes verify protein expression, while IgG serves as a negative control. (F) Co-IP experiments validating specific interactions between FLAG-ZRF1 and candidate interactors. Left: Interaction between FLAG-ZRF1 and SERBP1-GFP, and reciprocal IP using SERBP1-GFP and FLAG-ZRF1. Middle left: Co-IP showing binding between FLAG-ZRF1 and HABP4-GFP. Middle right: FLAG-ZRF1 precipitates with DDX17-Myc, DDX21-V5, and DDX5-GFP, confirming association. Bottom: GFP-tagged ZRF1 co-precipitates with FLAG-AGO1.

**Supplemental Figure 6:**
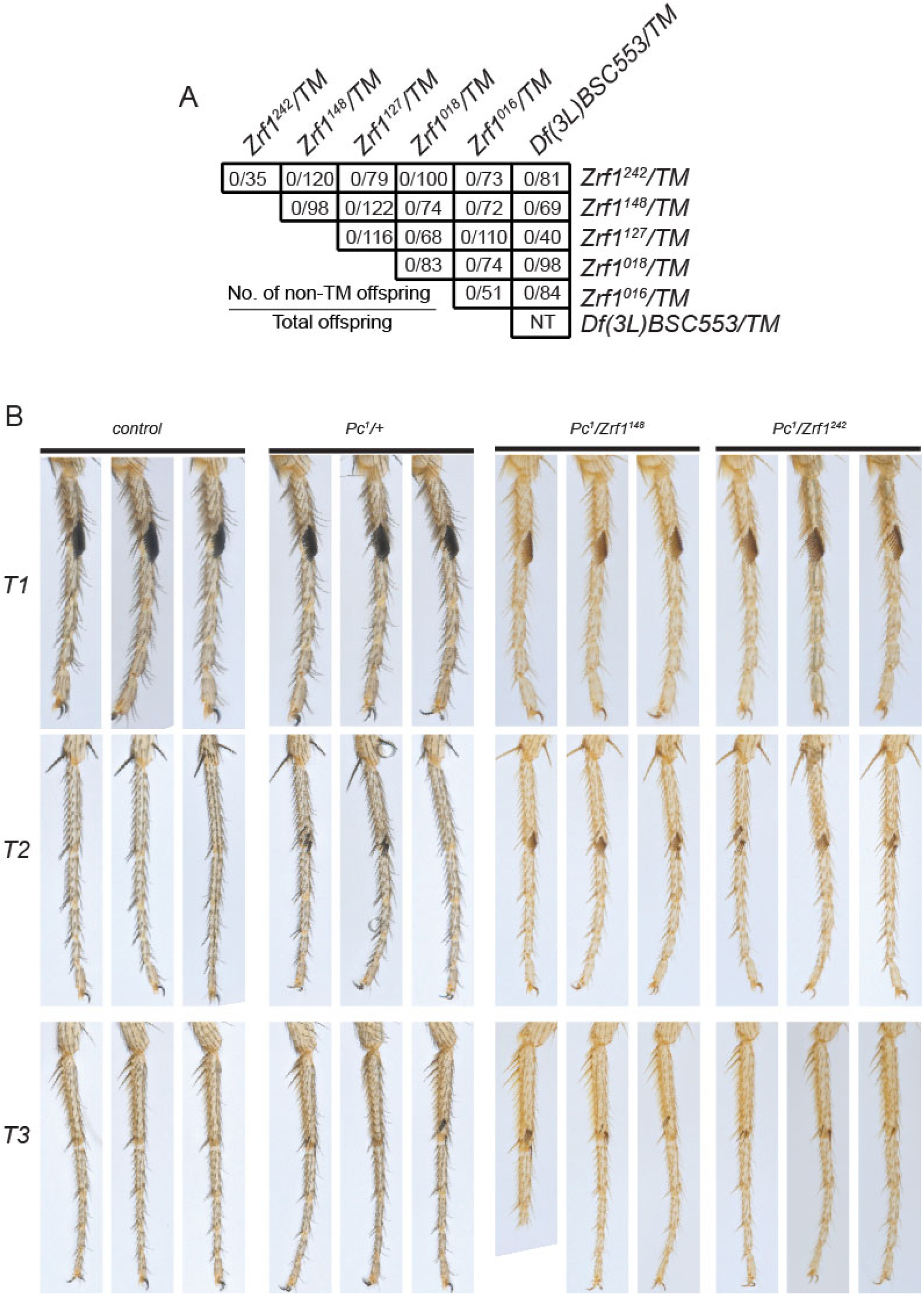
Complementation Testing and Genetic Interaction Between *Zrf1* and *Pc*. (A) Table showing the results of complementation testing between different *Zrf1* alleles and a deficiency line (*Df(3L)BSC553*) that covers the *Zrf1* locus. The number of non-TM (non-Tubby marker) offspring is listed against the total number of offspring, indicating whether the tested *Zrf1* alleles fail to complement each other or the deficiency. The absence of non-TM offspring suggests that the alleles are functionally compromised and do not complement, consistent with loss-of-function mutations. (B) Images of male forelegs (T1), midlegs (T2), and hindlegs (T3) from control, *Pc^1^/+* heterozygous, and double heterozygous *Pc^1^/Zrf1* mutants (*Zrf1^148^, Zrf1^2^*^42^). Control males have normal sex combs restricted to T1. *Pc*^1^/+ heterozygotes display ectopic sex combs on T2 and T3, and this phenotype is enhanced in *Pc^1^/Zrf1* double heterozygotes, indicating a genetic interaction between *Pc* and *Zrf1*.

**Supplemental Figure 7:**
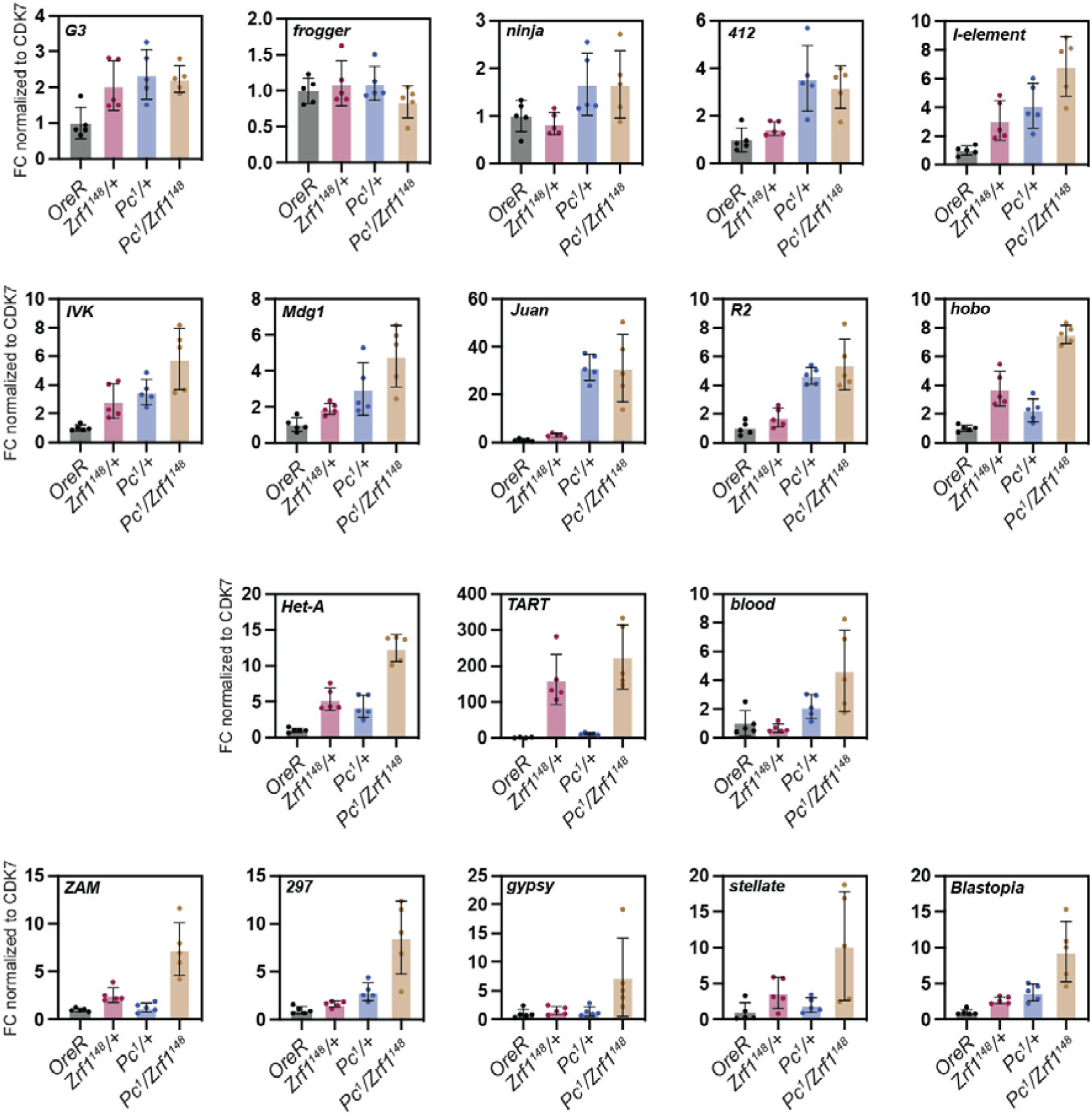
Quantitative Analysis of Retrotransposable Element (TE) Expression in *Zrf1* and *Pc* Mutants. Graphs displaying the fold change (FC) in expression of various retrotransposable elements (TEs) normalized to *CDK7* in different genetic backgrounds: *OreR* (wild-type control), *Zrf1^148^/+*, *Pc^1^/+*, and double heterozygous *Pc^1^/Zrf1^148^*. TEs analyzed include *G3, frogger, ninja, 412, I-element, IVK, Mdg1, Juan, R2, hobo, Het-A, TART, blood, ZAM, 297, gypsy, stellate*, and *Blastopia*. Each bar graph shows relative TE expression, indicating how the presence of *Zrf1* and *Pc* mutations affects TE regulation. Error bars represent standard deviation, and individual data points indicate biological replicates.

